# Reverse genetic screen identifies malaria parasite genes required for gametocyte-to-sporozoite development in its mosquito host

**DOI:** 10.1101/2023.03.14.532540

**Authors:** Chiamaka Valerie Ukegbu, Ana Rita Gomes, Maria Giorgalli, Melina Campos, Alexander J. Bailey, Tanguy Rene Balthazar Besson, Oliver Billker, Dina Vlachou, George K. Christophides

**Affiliations:** Department of Life Sciences, Imperial College London, London, SW7 2AZ, UK; Wellcome Trust Sanger Institute, Wellcome Trust Genome Campus, Hinxton, CB10 1SA, UK; LPHI, CNRS, University of Montpellier, 34095 Montpellier, France; Department of Pathology, Microbiology and Immunology, School of Veterinary Medicine, University of California Davis, Davis, CA 95616, USA; Department of Molecular Biology, Umea University, Umea, 90187, Sweden; Laboratory for Molecular Infection Medicine Sweden, Umea, 90187, Sweden

## Abstract

Malaria remains one of the most devastating infectious diseases. Reverse genetic screens offer a powerful approach to identify genes and molecular processes governing malaria parasite biology. However, sexual reproduction and complex regulation of gene expression and genotype-phenotype associations in the mosquito have hampered the development of screens in this key part of the parasite lifecycle. We designed a genetic approach in the rodent parasite *Plasmodium berghei*, which in conjunction with barcode sequencing allowed us to overcome the fertilization roadblock and screen for gametocyte-expressed genes required for parasite infection of the mosquito *Anopheles coluzzii*. The results confirmed previous findings, validating our approach for scaling up, and identified new genes required for ookinete motility and mosquito midgut infection and for sporozoite development and oocyst egress and salivary gland infection. Our findings can assist efforts to study malaria transmission biology and develop new interventions to control disease transmission.

## Introduction

Enhanced vector control has reduced the number of malaria cases and, together with effective medicines and improved health care, helped reduce the number of malaria-related deaths. Nevertheless, the latest reports indicate that these measures alone cannot bring further impact on disease control. Mosquito resistance to insecticides used in bed-nets and indoor residual spraying has increased, and mosquito biting and resting behavior has changed partly because of these measures. Since the only antimalarial vaccine licensed to date (RTS, S) is predicted not to have a universal game-changing impact, additional tools are needed, especially those that target disease transmission.

Malaria is caused by the protozoan parasite *Plasmodium* that is transmitted between humans through bites of *Anopheles* mosquitoes. Transmission commences when a female mosquito ingests haploid *Plasmodium* gametocytes that inside the mosquito midgut rapidly produce gametes that produce zygotes upon fertilization. Whilst undergoing meiosis, a diploid and then tetraploid zygote becomes a motile ookinete and traverses the midgut epithelium. During midgut traversal, the majority of ookinetes are eliminated by the mosquito immune system. In the midgut sub-epithelial space, an ookinete differentiates into a replicative oocyst where thousands of haploid sporozoites develop within a period of about 2 weeks through endomitotic replication and budding. Following egress from the oocyst, the sporozoites migrate to the salivary glands from where they are transmitted to another human upon a next mosquito bite.

The relationship between gene expression and protein function throughout *Plasmodium* development is multifaceted. Within the mosquito, parasite developmental transitions coincide with notable changes in transcriptome repertoires (Akinosoglou et al., 2015; Ukegbu et al., 2015), facilitated by transcription factors of the Apetala 2 (AP2) family (Modrzynska et al., 2017; Yuda et al., 2009). However, post-transcriptional regulation plays a pivotal role in *Plasmodium* transmission biology (Hall et al., 2005; Lasonder et al., 2016), exemplified by the synthesis and storage of transcripts in the female gametocyte, which are released for translation after fertilization (Guerreiro et al., 2014; Mair et al., 2006; Mair et al., 2010). Amongst these is the AP2-O transcription factor that controls *de novo* gene expression in the zygote and ookinete. Furthermore, some proteins produced in ookinetes are transported into an organelle exclusive to ookinetes and young oocysts, called crystalloid, and appear to function later during oocyst development (Dessens et al., 2021).

The spatiotemporal mismatch between parasite gene expression and protein function together with increased ploidy in the zygote and ookinete and endomitosis in the oocyst have hindered efforts to develop genetic screens to study mosquito infection and disease transmission. Efficient tools combining the scalability of signature tagged mutagenesis (STM) and the throughput of barcode sequencing have enabled high throughput genetic screens in the haploid asexual blood stages (ABS) of the rodent parasite *Plasmodium berghei* (Bushell et al., 2017; Gomes et al., 2015). However, cross-fertilization between mutants in the mosquito blood bolus limits the utility of barcoded mutants for identifying gene functions before products of meiosis and endomitotic replication cycles segregate during sporogony (Stanway et al., 2019), highlighting the need for more elaborate genetic designs.

Here, we present a new reverse genetics screen design that enables the study of genes involved in the gametocyte-to-sporozoite development. Guided by our earlier discovery that the male *P. berghei* genome is largely inactive in the first 32 hours in the mosquito (Ukegbu et al., 2015), our approach involves STM in parasites producing only female gametes that are then crossed to parasites producing only male gametes, leading to zygotes that carry female null and male wildtype alleles. Apart from preventing barcodes from being transmitted through male gametocytes, this approach precludes the generation of double mutants that could reduce the analytical power of the screen when co-inherited mutations have strong and/or interacting phenotypes. We first validated the design by confirming phenotypes of previously studied genes and concluded that this is a powerful strategy for scaling up the rate at which functions could be assigned to genes transcriptionally enriched in gametocytes. We then characterized three such unstudied genes identified by the screen, all encoding putative transmembrane proteins, and showed that *STONES* encodes a protein associated with the Ookinete Extrados Site (OES) and required for ookinete motility, while *CRYSP* and *CRONE* encode crystalloid proteins required for sporozoite formation and oocyst egress and/or salivary gland infection, respectively. Last, we investigated two genes not detected by the screen but identified in our published and unpublished data as important for transmission. We confirmed that *PIMMS57*, which we previously showed to encode a protein required for oocyst development (Ukegbu et al., 2021), and *ROVER*, which encodes a previously uncharacterised protein essential for ookinete motility associated with vesicle trafficking, have knockout phenotypes that are fully rescued by the male wildtype alleles. This is consistent with the predicted shortcoming of the screen to reveal recessive phenotypes in diploid cells, further proving the validity of our design for the discovery of genes and processes important for *Plasmodium* transmission biology.

## Results and discussion

### Identification of gametocyte-enriched transcripts *in vivo*

To identify gametocyte-enriched genes *in vivo* in the mosquito midgut, which we could then include in the screen, we infected *A. coluzzii* (previously *A. gambiae* M form) with the *P. berghei* ANKA 2.34 or ANKA 2.33 (non-gametocyte producing) lines, and used RNA isolated from the mosquito midguts 1 hour post blood feeding (pbf) in competitive hybridizations of a *P. berghei* oligonucleotide microarray (Akinosoglou et al., 2015). Three replicate infections and respective hybridizations were performed that allowed us to examine the expression of 3,428 genes, after excluding probes ambiguously mapping to the genome. Data analysis identified 189 transcripts with significant >1.6-fold enrichment in ANKA 2.34 compared to ANKA 2.33 (**Table S1**). The expression of 109 of these genes was previously shown to be affected in parasites lacking the DEAD-box RNA helicase DOZI (development of zygote inhibited) which is essential for translational mRNA repression in female gametocytes (Mair et al., 2010); DOZI itself was among these 109 genes. Comparison with data obtained from single cell RNA sequencing of gametocytes (Howick et al., 2019) showed that 74% of the upregulated genes (139 genes) are highly expressed in gametocytes with 83% of these genes (116 genes) being enriched in female gametocytes. The full analysis is presented in the **Supplemental Information** and in **Figure S1**.

### STM screen optimization

We created a library of barcoded knockout *Plasmo*GEM vectors targeting gametocyte-enriched genes identified as above, as well as control genes of known or no phenotypes. In this initial study, we pooled 27 of these vectors (**Table S2**) targeting 13 previously characterized genes (including PBANKA_0720900), 6 genes encoding 6-cys-containing proteins (including P36 detected in our transcriptomic study), and 5 uncharacterized genes detected in our transcriptomic study and previously found to be translationally repressed by DOZI (Mair et al., 2010): PBANKA_0413500 (*STONES*), PBANKA_1338100 (*CRYSP*), PBANKA_1353800 (*ROVER*), PBANKA_0810700 (*SPM1*) and PBANKA_0720900 (*CRONE*). The pool also included 4 epitope-tagging *Plasmo*GEM vectors previously shown not to impose any fitness cost upon ABS development (Bushell et al., 2017; Gomes et al., 2015). Thus, the total number of *Plasmo*GEM vectors in the pool was 31.

STM in wildtype (*wt*) *P. berghei* has been shown to be effective in ABS screens (Bushell et al., 2017; Gomes et al., 2015) but when viable mutants were transmitted through the life cycle, the power to detect phenotypes specific to one sex or during the diploid phase of the life cycle was limited until products of meiosis had segregated during sporogony (Stanway et al., 2019). We sought to overcome the latter by mutagenesis in a female-donor line that would be then crossed to a male-only donor line leading to unidirectional crosses and no double mutants (**Figure 1A**). In this design, complementation by the *wt* male alleles is addressed by a previous finding that the male genome is largely silent during the first 32 hours after fertilization (Ukegbu et al., 2015), except for genes specifically expressed in the zygote/ookinete. Thus, our design allows to screen for phenotypes in mosquito stages associated with gene expression in the female gametocyte, mostly caused by maternal transcript or protein deposition, or with constitutive gene expression.

**Figure 1.**
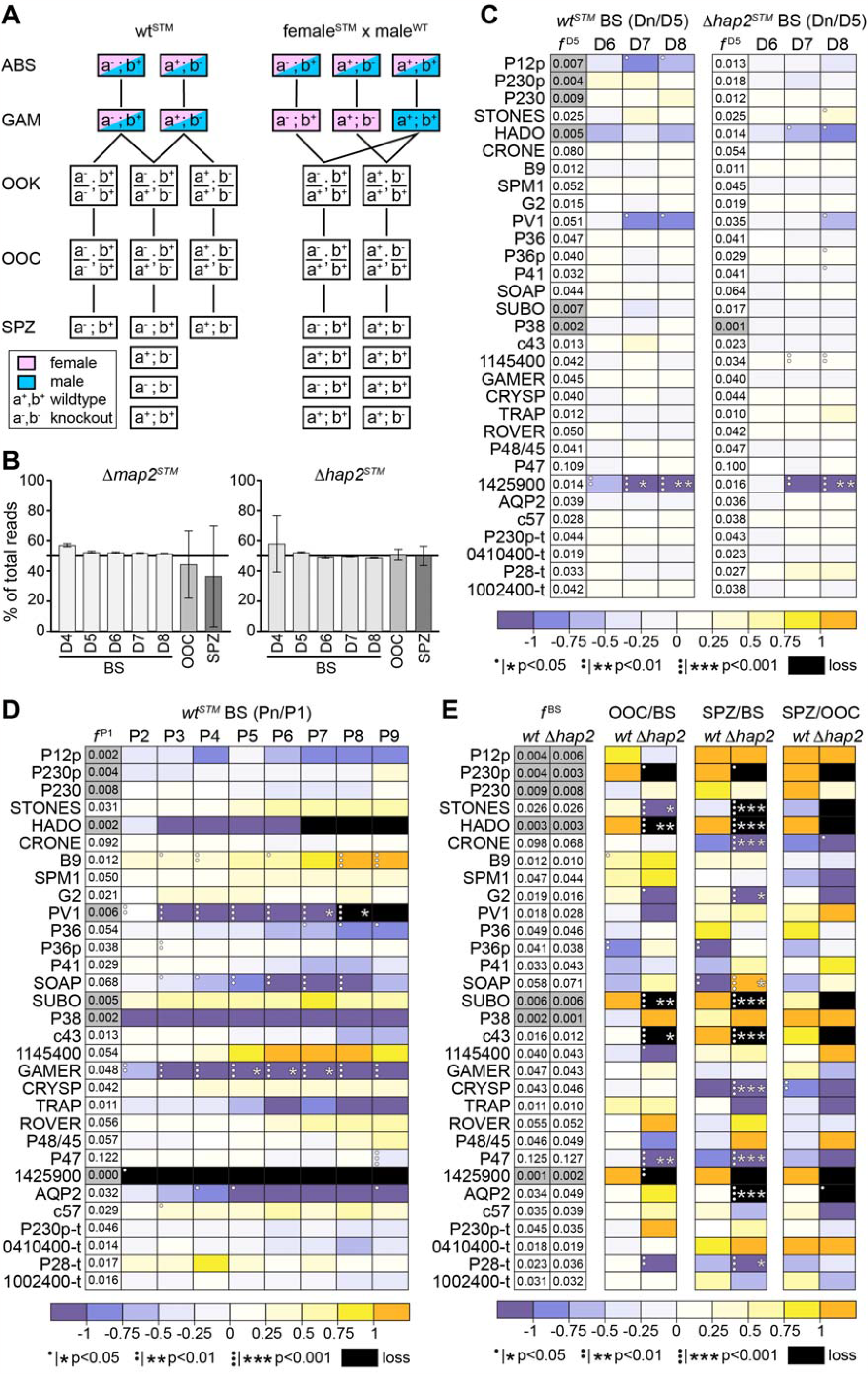
Design and results of the STM screen. **(A)** Parasite genotypes after mutagenesis in *wt* parasites (wt^STM^) or in a female-donor line (female^STM^; *e*.*g*., *Δmap2*^*STM*^ and *Δhap2*^*STM*^) crossed to a *wt* male-donor line (male^WT^; ; *e*.*g*., *Δnek4*^*WT*^). An example of two mutated loci (a and b) is presented. ABS, asexual blood stages; GAM, gametocytes and gametes; OOK, ookinetes; OOC, oocysts; SPZ, sporozoites. **(B)** Growth dynamics of the *Δmap2* and *Δhap2* lines in mouse blood stages (BS) at days 4-8 (D4-D8) after transfection with the STM pool and in oocyst (OOC) and salivary gland sporozoite (SPZ) stages after crossing each line to *Δnek4* prior to infection of *A. coluzzii* mosquitoes. The percentage of map2 and hap2 barcode counts in the total barcode counts is shown. Whiskers show standard error of mean. **(C)** Fitness of blood stage (BS) STM mutants in the *wt* (left) and *Δhap2* (right) genetic backgrounds shown as relative barcode abundance (ratio of each to total barcode counts) at days 5-8 (Dn) to day 5 (D5) post mouse infection. *f* ^D5^ is the frequency of each barcode in every 1000 barcodes at the D5 sample. **(D)** Stability of STM mutants in the *wt* genetic background throughout 9 successive mouse-to-mouse passages (P1-P9) shown as relative barcode abundance (ratio of each to total barcode counts) in each passage (Pn) to the first passage (P1). *f* ^P1^ is the frequency of each barcode in every 1000 barcodes in the P1 sample. **(E)** Developmental progression of STM mutants in *wt* or *Δhap2* genetic backgrounds inside *A. coluzzii* mosquitoes, shown as relative barcode abundance (ratio of each to total barcode counts) in oocysts (OOC) and sporozoites (SPZ) to blood stages (BS), or SPZ to OOC. *f* ^BS^ is the frequency of each barcode in every 1000 barcodes in the blood stage sample. For panels (C), (D) and (E), differences in abundance are color-coded as shown at the bottom of each panel, while grey-shaded boxes denote barcodes with starting frequencies less than 0.01. Statistical analysis is done with a student’s t test, and p values are shown as dots prior to multiple testing correction and as stars after multiple testing correction. Black boxes indicate zero or near zero count ratios.

After initial experiments, we chose to test two female-donor lines: *Δmap2* and *Δhap2*. Absence of the mitogen-activated protein kinase MAP2 inhibits male gamete exflagellation by impairing cytokinesis and axoneme motility (Billker et al., 1997; Tewari et al., 2005; Tewari et al., 2010), while absence of the male-specific membrane fusogen HAP2 renders the male gametes unable to fuse with the female gametes (Hirai et al., 2008; Liu et al., 2008; Mori et al., 2010). We used barcoded *Plasmo*GEM vectors and a negative selection strategy (Braks et al., 2006) to generate *Δmap2* and *Δhap2* selection marker free lines (**Figure S2A**). After transfection with the pool of barcoded vectors, each of the transfectant lines was used to coinfect mice together with a barcode-free male-donor *Δnek4* line (Reininger et al., 2005; Reininger et al., 2009), leading to fertilization of the mutant female gametes by *wt* male gametes once inside the *A. coluzzii* midguts. We sampled blood from mice infected with each of the transfected lines at days 4-8 post infection (pi), and mosquito midguts and salivary glands at days 12 and 21 pbf. Thus, each transfected parasite was expected to carry two barcodes: one that identifies the genetic background (*Δmap2* or *Δhap2*), expected to account for 50% of the total barcode counts, and one from the 31-vector pool. The abundance of each of the barcodes and its change over time would allow us to track the growth dynamics of the mutant populations.

The results showed that from day-5 after mouse infection onward, once drug selection eliminated parasites that had not integrated a barcoded vector, the *map2* and *hap2* barcode counts were stabilized to about 50% of the total, as expected, with very little variability observed between replicates (**Figure 1B**). However, in oocysts and salivary gland sporozoites only the *hap2* barcodes were consistent between replicates and at about 50% of the total, while the *map2* barcodes were highly variable and less than 50% of the total. This indicated that the *Δmap2* line may exhibit defective sporozoite development and thus would not be suitable for our screen; hence, we continued with the *Δhap2* female-donor line.

### Fitness and stability of mutants in the mouse host

We assessed the fitness of each STM mutant parasite in the mouse host, expressed as relative barcode abundance (ratio of counts of each barcode to all barcodes) in days 5-8 (Dn) compared to day 5 (D5) pi (**Figure 1C**). This was done both in the validated *Δmap2* and also in a *wt* genetic backgrounds, which would allow us to identify and exclude any genetic interactions between the pooled genes and *HAP2* (Fang et al., 2018). As expected, parasites lacking the gene *PBANKA_*1425900 that was previously reported to have a reduced ABS growth rate (Bushell et al., 2017), which was used here as a control, exhibited drastically reduced fitness in both the *wt* and *Δhap2* genetic backgrounds, which remained statistically significant after false discovery rate (FDR) correction at days 7 and 8 pi. The abundance of all other mutants did not change significantly throughout the course of the infection.

Because *P. berghei* lines must be serially maintained in rodents either through direct blood passages until a fully replicated experiment is finished or with intermediate freezing and thawing of the infected blood, although this presents an additional stress for the parasite, we wanted to examine the stability of the mutant parasite population during this process. We investigated the stability of the mutant population in the *wt* genetic background over 9 successive mouse-to-mouse passages (P1-9), by comparing the relative barcode abundance in each passage with that in the first infected mouse (**Figure 1D**). Three mutants were found to dropout at different points of this experiment due to either reduced ABS growth rate or very low abundance in the pool following transfection: *PBANKA_1425900* (P2), *HADO* (P7) and *PV1* (P8). *HADO* encodes a putative magnesium phosphatase important for ookinete development, possibly by regulating the actin dynamics (Akinosoglou et al., 2015), and *PV1* encodes a homolog of an essential protein of the *P. falciparum* parasitophorous vacuole (Chu et al., 2011), likely an accessory of the translocon protein complex PTEX (Morita et al., 2018). Another mutant that started with a relatively high abundance (0.048) but was significantly depleted from the parasite population from P5 onward was *GAMER* (**Figure 1D**). While *GAMER* is shown to be important for male gamete release (Akinosoglou et al., 2015), it is also highly expressed in ABS where it may have a non-essential or redundant role that eventually caused the loss of the mutant from the population due to fitness cost. From these results, we concluded that, albeit serial mouse passages having little or no impact on screening genes expressed in gametocytes onwards, the parasite population should be better kept at a low passage stage to avoid the loss of mutants with low starting frequency.

### Oocyst and salivary gland sporozoite development in the mosquito host

The ability of the *P. berghei Plasmo*GEM mutants to infect *A. coluzzii* and develop to oocysts which would then produce sporozoites that migrate and infect the salivary glands was examined at days 12 and 21 pbf, respectively. Mutagenesis in both the *wt* and *Δhap2* genetic backgrounds was investigated; the *Δhap2* female-donor mutants being crossed to the male-donor *Δnek4* through mouse coinfections. The ratio of normalized barcode counts in oocysts and salivary gland sporozoites to mouse blood stages prior to mosquito blood feeding was calculated for every mutant in four independent replicates. Data analysis revealed a stark difference between the two genetic backgrounds: while several *Δhap2 Plasmo*GEM mutants appeared to be significantly affected or completely dropout from the screen at either or both of these stages, no significant effect was observed for any mutant in the *wt* background likely due to complementation with a functional allele upon fertilization (**Figure 1E**).

Three mutants (*Δhado, Δsubo* and *Δcpimms43*, aka *Δc43*) completely dropped-out and another two mutants (*Δsto* and *Δp47*) were drastically depleted from the parasite population at the oocyst stage (**Figure 1E**). *Δsto* that carries a disruption of a previously unstudied gene, *STONES*, completely dropped-out from the population in the salivary gland sporozoite stage. As said above, *HADO* is required for ookinete development and transformation to oocyst, with a possible role in midgut invasion (Akinosoglou et al., 2015). Both *PIMMS43* and *P47* (van Dijk 2010) are essential for ookinete protection from mosquito complement reactions upon midgut traversal (Ukegbu et al., 2020; Ukegbu et al., 2017b). P47, a 6-cys domain protein also found on the surface of female gametocytes, is additionally important for gamete fertilization (Ukegbu et al., 2017b; van Dijk et al., 2010). *SUBO* (aka *PIMMS2*) encodes an ookinete-specific subtilisin-like protein required for ookinete traversal of the midgut epithelium, possibly being involved in epithelial cell egress (Ukegbu et al., 2017a). Two additional mutants, *ΔPBANKA_1425900* and *Δp230p*, also dropped-out from the screen at this stage. P230p is a paralogue of P230, a 6-cys domain protein involved in fertilization, and its *P. falciparum* ortholog is shown to be important for early parasite development in the mosquito (Marin-Mogollon et al., 2018).

Five mutants showed reduced salivary gland sporozoite albeit normal oocyst growth. *Δaqp2* that carries a disruption of *AQP2*, a gene encoding a protein with high similarity to aquaglyceroporins, completely dropped-out from the salivary gland sporozoite population, while the relative abundances of *Δcro* (disruption of *CRONE*), *Δcry* (disruption of *CRYSP*), *Δg2* (disruption of *G2*) and *p28-t* (tagged *P28*) were significantly reduced (**Figure 1E**). *CRONE* (PBANKA_0720900), which encodes a 265 amino acid protein with an N-terminal signal peptide, has been previously shown to be expressed in gametocytes, where it is translationally repressed by DOZI, and translated in ookinetes to a protein localized in the crystalloids and required for sporozoite production in the oocyst (Guerreiro et al., 2014). *CRYSP* (PBANKA_1338100) encodes a previously unstudied 263 amino acid protein with 3 transmembrane domains. The ookinete and sporozoite protein G2 (glycine at position 2) has been previously shown to localize in the cortical subpellicular network of these zoite stages and be essential for their morphogenesis (Tremp et al., 2013). Finally, *p28-t* is designed to produce a C-terminally 3x human influenza hemagglutinin (3xHA) tagged version of the ookinete P28 GPI-anchored protein, a known target of transmission blocking vaccines (Tomas et al., 2001), and was used here as a negative control. Although P28 is known to have a redundant function, it appears that its tagging leads to a malfunctional protein that affects parasite development in the vector. Indeed, a drastic reduction of *p28-t* was already seen in the oocyst stage although this was not statistically significant following FDR correction.

### Detailed phenotypic analysis of single mutant parasites

We selected for further analysis, three genes with strong phenotypes, *STONES, CRYSP* and *CRONE*, as well as two genes for which the screen revealed no phenotype, *ROVER* and *SPM1*; an independent preliminary study had indicated that *ROVER* may be involved in infection, while *SPM1* was used as a control. *In silico* analysis of each of the predicted proteins and amino acid sequence alignments of selected orthologs from other *Plasmodium* species are presented in **Figures S3-S7**, respectively. Briefly, in addition to *CRYSP* and *CRONE* described earlier: *STONES* encodes a 1,037 amino acid protein with 14 transmembrane domains and coupled N-terminal LIS1 homology (LisH) motifs; *ROVER* encodes a 367 amino acid protein with no predicted domains; and *SPM1* encodes a 329 amino acid putative subpellicular microtubule (SPM) protein predicted to contain a microtubule associated protein 6 (MAP6) domain.

We used the *Plasmo*GEM disruption vectors for *STONES* and *CRYSP* **(Figure S2B)** and conventional disruption vectors for *CRONE, ROVER* and *SPM1* **(Figure S2C)** to generate clonal *P. berghei* mutants in the *c507* GFP-expressing *wt* line. Integration of the disruption cassettes and gene deletion in the clonal *Δsto, Δcry, Δcro, Δrov* (*ROVER*) and *Δspm1* parasite lines was confirmed by PCR **(Figure S2B**,**C)**.

For all mutant parasites, male gametogenesis, determined as the number of *in vitro* recorded exflagellation events per the number of male gametocytes, was comparable to that of the parental *c507 wt* control **(Figure 2A)**. Ookinete conversion rates, *i*.*e*., the ratio of ookinetes to female gametocytes counts were also not significantly different from the control for all the mutants **(Figure 2B)**. Next, we examined the ability of the mutant parasites to form oocysts in *A. coluzzii*, following feeding of mosquitoes on mice infected with each of these mutants or the *c507 wt* control line. Oocyst counts were determined 8 days pbf. *Δcry, Δcro* and *Δspm1* mutants produced oocysts that were not significantly different in number from the control **(Figure 2C, Table S3)**. However, a 99% decrease of mean oocyst numbers was observed for *Δsto* and *Δrov* mutants, with *Δrov* showing a maximum of only one oocyst in some midguts.

**Figure 2.**
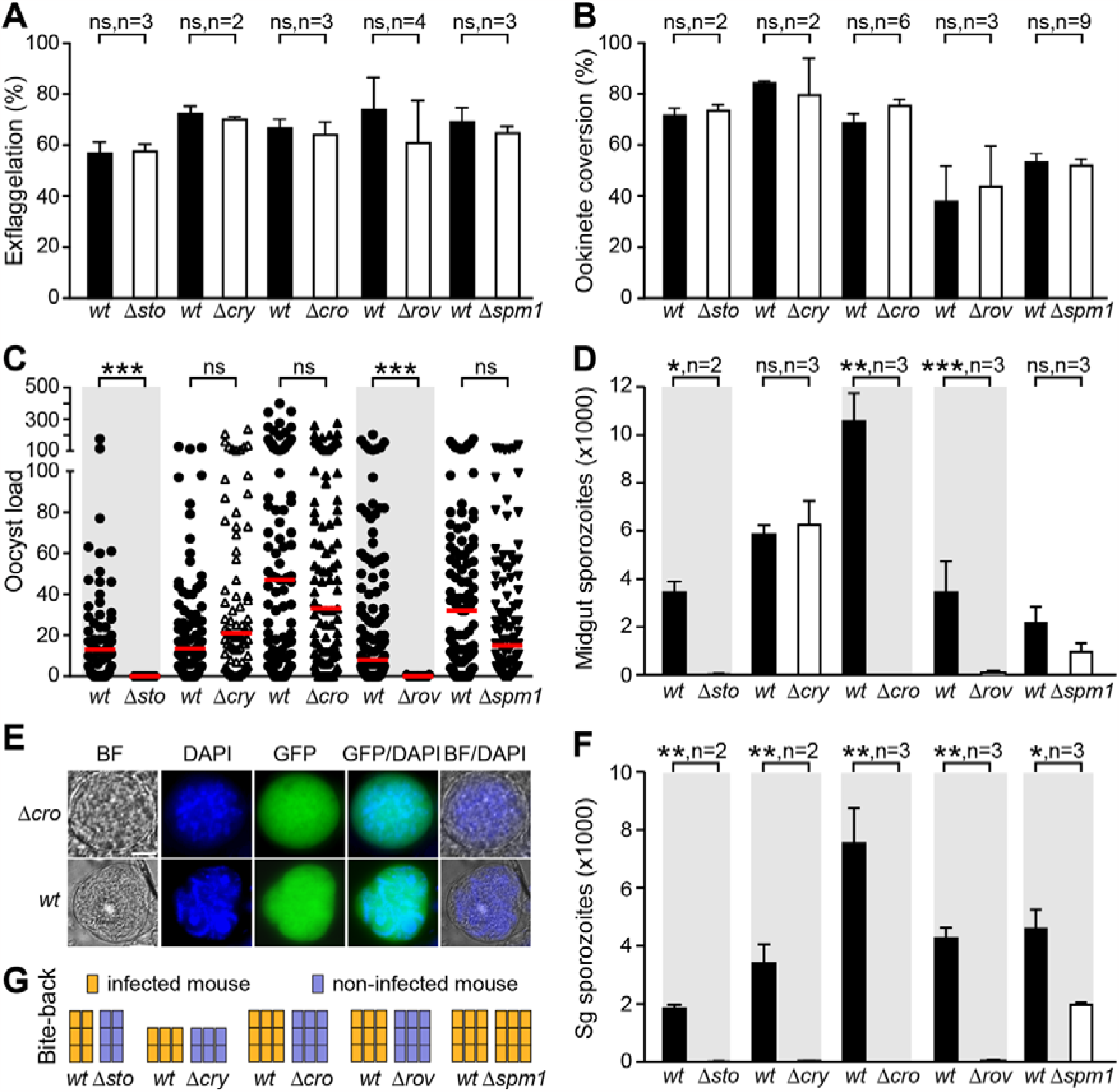
Phenotypic characterization of *P. berghei* knockout mutant parasites. **(A)** *In vitro* exflagellation capacity of mutant parasites shown as percentage of exflagellation of male gametocytes compared to *c507 wt* controls. Whiskers show standard error of mean (SEM). Statistical analysis is done with the student’s t test. ns; not significant, n; number of biological replicates. **(B)** *In vitro* female gamete to ookinete conversion of mutant parasites compared to *c507 wt* controls. Whiskers show SEM. Statistical analysis is done with the student’s t test. ns, not significant; n, number of biological replicates. **(C)** Oocyst load in *A. coluzzii* midguts of mutant parasites compared to the *c507 wt* parasite at day 8 pbf. Red lines show median. Statistical tests were performed using Mann-Whitney. ***, P<0.0001; ns, not significant. Gray shading highlight statistically significant differences. **(D)** Number of sporozoites in *A. coluzzii* midguts of mutant parasites compared to *c507 wt* controls. Whiskers show SEM. Statistical analysis is done with the student’s t test. *, P<0.05; **, P<0.001; ***, P<0.0001; ns, not significant; n, number of biological replicates. Gray shading highlight statistically significant differences. **(E)** Fluorescence microscopy images of GFP expressing *Δcro* oocysts compared to *c507 wt* controls at day 15 pbf in *A. coluzzii* midguts. Midguts were stained with DAPI (blue). BF; brightfield. Scale bar is 5 μM. **(F)** Number of sporozoites in *A. coluzzii* salivary glands (sg) of mutant parasites compared to *c507 wt* controls. Whiskers show SEM. Statistical analysis is done with the student’s t test. *, P<0.05; **, P<0.001; n, number of biological replicates. Gray shading highlight statistically significant differences. **(G)** Mouse infection from bite-back experiments of mosquitoes infected with mutant parasites or *c507 wt* controls. Each mouse is shown as a rectangle and columns indicate independent biological replicates. Infected mice are shown with yellow color and non-infected mice are shown with blue color.

Whilst the *STONES* phenotype was consistent with that obtained from the screen, the *ROVER* phenotype was unexpected and could only be justified by re-expression of the gene in the ookinetes and male *wt* allele rescue of the phenotype in the screen. We investigated this by crossing *Δrov* to either the female-donor *Δhap2* or the male-donor *Δnek4* followed by oocyst counting in *A. coluzzii* 8 days pbf on coinfected mice. The *Δc57* line that harbors a disruption of *PIMMS57* was also included in these assays, as the screen also failed to detect this gene that has been previously shown to be important for ookinete-to-oocyst transition (Ukegbu et al., 2021). The results confirmed that the oocyst-deficient phenotypes of both genes can be rescued by both the male and female *wt* alleles (**Figure S8, Table S4**), consistent with the expected limitation of the screen to reveal recessive phenotypes in diploid cells, after the *wt* allele introduced into the zygote by the microgamete is transcribed. *PIMMS57* is known to be specifically expressed in ookinetes (Ukegbu et al., 2021), likely by both parental alleles, and the results suggest that the gametocyte-enriched *ROVER* gene is also expressed in ookinetes and that this expression is important for its function.

Next, we assessed the capacity of mutant parasites to produce sporozoites that can migrate to the salivary glands, by counting midgut (oocyst) and salivary gland sporozoites 15 and 21 days pbf, respectively. Consistent with a defective ookinete-to-oocyst transition, very few *Δsto* (10±7) and *Δrov* (61±11) oocyst sporozoites were detected (**Figure 2D, Table S5**), while the *Δcro* oocysts were also devoid of sporozoites consistent with what has been reported previously (Guerreiro et al., 2014). As the screen did not detect any significant reduction in oocyst *CRONE* barcode counts, we carried out microscopy on mature *Δcro* oocysts 15 days pbf to further investigate this phenotype. The results revealed that *Δcro* oocysts had large nuclei filled with DNA but these were highly disorganized, unlike *wt* oocysts that exhibited highly organized nuclei with haploid sporozoites budding off from the sporoblastoid body **(Figure 2E)**. These data indicated normal DNA replication in *Δcro* oocysts, but defective sporozoite formation and budding, further validating our genetic screen. Finally, the numbers of *Δcry* and *Δspm1* midgut sporozoites were not significantly different from those of *wt c507* controls, also corroborating the results obtained from the screen.

These results were also reflected in the salivary gland sporozoite counts for *Δsto, Δcro* and *Δrov*, which ranged from very few to none (**Figures 2F, Table S5**). Importantly, and consistent with the results of the screen, none of the thousands of *Δcry* oocyst sporozoites were capable of infecting the salivary glands, again corroborating the results of the screen. A statistically significant 57% reduction in sporozoite counts was detected for *Δspm1* suggesting that the effect seen in midgut sporozoites may also be true.

Finally, the ability of mutant parasites to transmit to the mouse host and infect RBCs was assessed using mosquito-to-mouse (C57/BL6 strain) bite-back infections 21 days pbf (**Figures 2G, Table S5**). As expected, no transmission and development of mouse parasitemia was detected for any of the *Δsto, Δcro, Δcry* and *Δrov* mutants, leading us to conclude that loss-of-function of the respective proteins leads to malaria transmission blockade. However, the reduction seen in *Δspm1* salivary gland sporozoites did not bear any impact on the capacity of mutant sporozoites to infect the mouse host, suggesting that SPM1 is dispensable for sporozoite development and transmission.

### STONES and ROVER are required for ookinete motility and mosquito midgut invasion

The endogenous *STONES, ROVER, CRYSP* and *CRONE* genes were tagged with C-terminal 3×HA tag via double crossover homologous recombination in the *c507* line, and the resulting transgenic lines were designated *stones::3xha, rover::3xha, crysp::3xha* and *crone::3xha*, respectively **(Figure S9)**.

We first analyzed *STONES* and *ROVER*, of which the disruption leads to defective ookinete phenotypes. The STONES::3×HA protein could not be detected at the predicted size of ∼ 125 kDa in Triton X-100 soluble extracts of blood stages, gametocytes or mature ookinetes. Instead, a band size of ∼ 16 kDa was detected predominantly in mature ookinetes and less in blood stages and gametocytes **(Figure 3A)**. However, in Triton X-100 insoluble extracts, a band of ∼ 80 kDa was specifically detected in mature ookinetes, with traces of it also seen in gametocytes. As the full-length protein was never detected in any of the extracts, these results suggest that STONES undergoes proteolytic processing, and that its C-terminal ∼80kDa fragment is embedded within the membrane owing to the multiple transmembrane domains. The ROVER::3xHA protein was detected only in mature ookinetes as 2 bands: the first at the expected size of ∼ 43 kDa, and the second, more predominant band at ∼25 kDa **(Figure 3B)**. This may indicate proteolytic cleavage of the protein.

**Figure 3.**
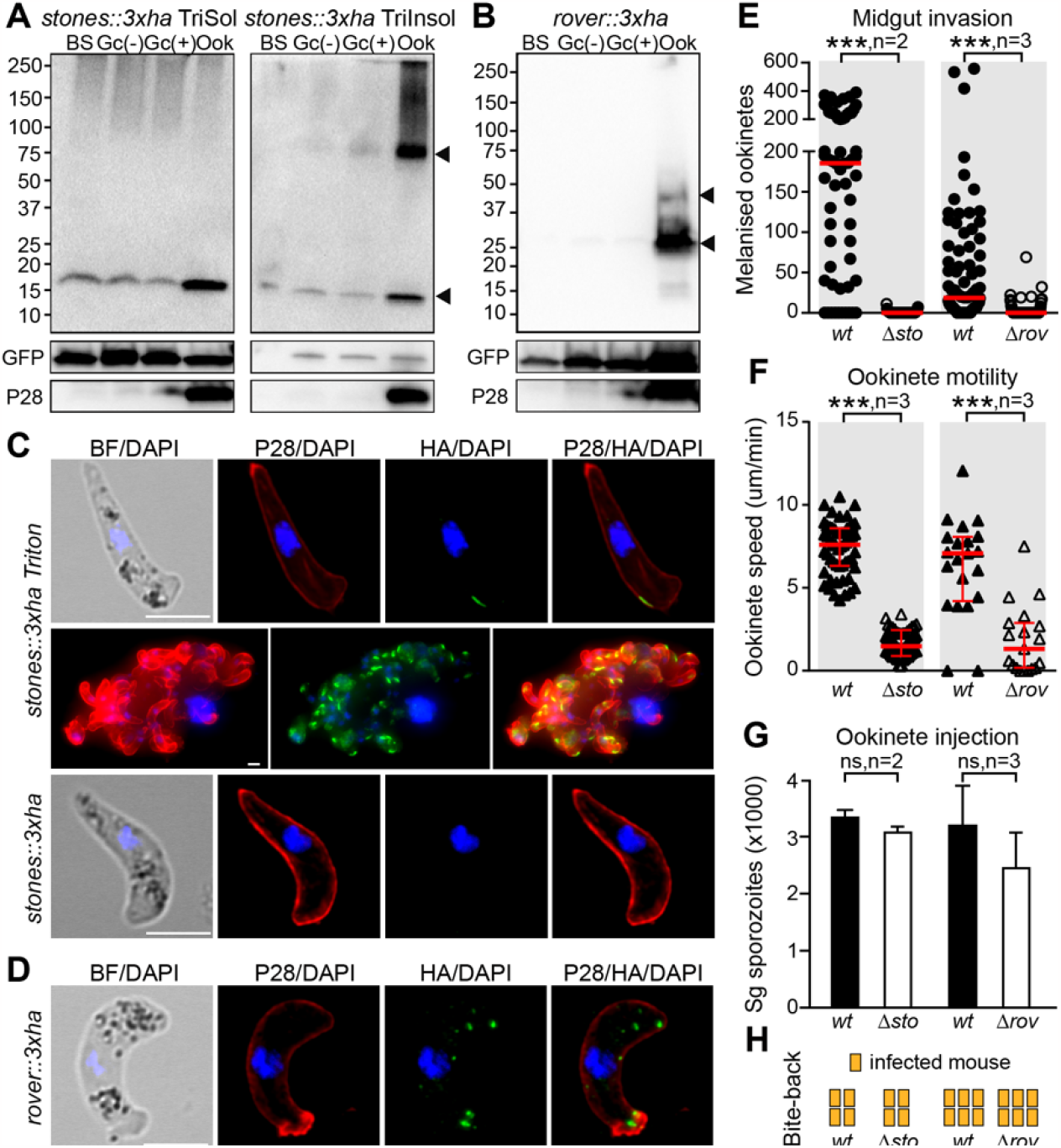
STONES and ROVER are involved in ookinete gliding motility. **(A)** Western blot analysis in the *stones::3xha* parasite line using an α-HA antibody under reducing conditions on Triton X-100 soluble (TriSol) and Triton X-100 insoluble (TriInsol) cell fraction. STONES::3HA specific signals are indicated with black arrowheads. GFP and P28 were used as loading and stage-specific controls, respectively. BS, mixed blood stages; Gc(-), inactivated and purified *in vitro* cultured gametocytes; Gc(+), activated and purified *in vitro* cultured gametocytes; Ook, purified *in vitro* cultured ookinetes. **(B)** Western blot analysis in the *rover::3xha* parasite line using an α-HA antibody under reducing conditions on whole cell lysates. ROVER::3XHA specific signals are indicated with black arrowheads. GFP and P28 were used as loading and stage-specific controls, respectively. BS, mixed blood stages; Gc(-), inactivated and purified *in vitro* cultured gametocytes; Gc(+), activated and purified *in vitro* cultured gametocytes; Ook, purified *in vitro* cultured ookinetes. **(C)** Immunofluorescence assays on *stones::3xha in vitro* cultured ookinetes that are Triton X-100 permeabilized (Triton; two top rows) and non-permeabilized (bottom row). Ookinetes are stained with α-HA (green) and α-P28 (red) antibodies. DNA was stained with DAPI (blue). BF; brightfield. Scale bar is 5 μM. **(D)** Immunofluorescence assays on *rover::3xha in vitro* cultured ookinetes. Ookinetes are stained with α-HA (green) and α-P28 (red) antibodies. DNA was stained with DAPI (blue). BF; brightfield. Scale bar is 5 μM. **(E)** Numbers of melanized ookinete in *CTL4* knock down *A. coluzzii* infected with the *Δsto, Δrov* and *c507 wt* control parasite lines. Red lines indicate median. Statistical analysis is done with the Mann-Whitney test. ***, P<0.0001; n, number of biological replicates. Gray shading highlight statistically significant differences. **(F)** Speed of *Δsto, Δrov* and *c507 wt* control *in vitro* cultured ookinetes measured with time-lapse microscopy, captured at 1 frame/5 sec for 10 min. Thick red lines indicate median and thinner red whiskers show SEM. Statistical analysis is done with the Mann-Whitney test. ***, P<0.0001; n, number of biological replicates. Gray shading highlight statistically significant differences. **(G)** Number of sporozoites in *A. coluzzii* salivary glands (sg) following injection of *Δsto, Δrov* and *c507 wt* control ookinetes cultured *in vitro* in the mosquito hemocoel. Whiskers show SEM. Statistical analysis is done with the student’s t test (unpaired two-tailed, equal variance). ns, not significant; n, number of biological replicates. **(H)** Mouse infection from bite-back experiments of mosquitoes infected with mutant *Δsto* or *Δrov* and *c507 wt* control parasites. Each mouse is shown as a rectangle and columns indicate independent biological replicates. Infected mice are shown with yellow color.

In immunofluorescence assays, STONES::3×HA was specifically detected at a distinctive membrane region located on the convex side of the mature ookinete, posterior to the apical structure **(Figure 3C)**. This region is critical for ookinete motility and has been termed Ookinete Extrados Site (OES) (Gao et al., 2018). In non-Triton X-100 treated mature ookinetes, no signal at the OES could be detected, suggesting that the N-terminal HA-tagged part of STONES is intracellular, which is consistent with its topology predictions. ROVER::3×HA was localized in discrete cytoplasmic spots of mature ookinetes, resembling exocytic vesicles, commonly but not always positioned toward the apical end and in proximity to the cell membrane **(Figure 3D)**.

The ookinete to oocyst defective phenotypes of the *Δsto* and *Δrov* parasites were further investigated in midgut invasion assays using a system we developed previously and involved infections of *A. coluzzii* silenced for *CTL4* (Ukegbu 2017). CTL4 is a key hemolymph regulator of melanization, and its silencing leads to readily melanized *P. berghei* ookinetes that have succeeded in invading the midgut epithelium and reached the sub-epithelial space (Osta et al., 2004). The results showed that both *Δsto* and *Δrov* ookinetes displayed a great defect in midgut invasion as the number of melanized ookinetes were significantly reduced compared to *wt* controls **(Figure 3E, Table S6)**.

Defective midgut invasion can be due to the inability of ookinetes to move, and we assessed this by measuring the forward speed of ookinetes on matrigel. The results confirmed that both *Δsto* and *Δrov* mutants exhibit strong motility defects which likely cause their decreased ability to traverse the midgut epithelium and form oocysts and sporozoites **(Figure 3F)**. To further examine this, *Δsto* and *Δrov* ookinetes were injected directly into the hemocoel to assess if the oocyst and sporozoite defective phenotypes could be rescued. Indeed, it has been previously shown that mosquito transmission of *P. berghei* mutants with ookinete motility defects can be rescued if midgut invasion is bypassed (Guttery et al., 2012). The result confirmed that this was the case for both *Δsto* and *Δrov*, as both the salivary gland sporozoite numbers and the ability of mutants for mouse transmission through bite-back were restored **(Figures 3G-H, Table S7)**.

Gliding motility is served by the glideosome, an actomyosin-based machinery located between the parasite plasma membrane (PPM) and the inner membrane complex (IMC) (Keeley and Soldati, 2004). Initiation of gliding in mature ookinetes coincides with the polarization of the PPM protein guanylate cyclase β (GCβ) to the OES (Gao et al., 2018). This leads to local elevation of cGMP levels and activation of cGMP-dependent protein kinase G (PKG) signaling that drives a series of events initiating gliding (Moon et al., 2009). Anchoring of GCβ and its cofactor CDC50A at the OES is facilitated by the IMC sub-compartment protein 1 (ISP1), which together with ISP3 interact with β-tubulin on the SPM, serving as tethers to maintain the proper SPM structure (Gao et al., 2018; Wang et al., 2020). However, it remains unclear what pulls GCβ/CDC50A to the OES in the first place, as ISP1 polarizes already at the zygote stage. Also, ISP1 is required for GCβ/CDC50A polarization in the majority but not in all the ookinetes, suggesting that additional proteins are involved in this process. The discovery of STONES (still ookinetes on the extrados site), a multi-transmembrane protein of the OES, may help shed new light into the mechanisms enabling this critical step in malaria transmission. The presence of the LisH motifs suggests that STONES contributes to the regulation of the SPM dynamics, either by mediating dimerization or by binding SPM directly. Whilst the STONES loss-of-function phenotype, cellular localization and predicted SPM association appear to be closely matching those of ISP1, our data cannot clarify whether STONES, like ISP1, is integral to the IMC or the PPM.

Ookinetes lack rhoptries and dense granules, and most of the proteins important for gliding motility and midgut invasion are trafficked to the membrane or extracellularly through the micronemes. These are specialized secretory organelles that are synthesized *de novo* in the Golgi and translocate apically using filamentous connections with the SPM (Schrevel et al., 2008; Sinden, 1999). Likewise, the ookinete IMC is thought to be formed *de novo* starting at the apical pole, most likely from Golgi-derived vesicles (Bannister et al., 2000), and observations in *Toxoplasma gondii* suggest that IMC recycling also happens (Ouologuem and Roos, 2014). The cellular localization of ROVER (roaming’s over) indicates an association with such vesicular structures. As the protein lacks a signal peptide and is never seen distributed across the membrane, it is suggestive that it acts as a cytoplasmic adaptor involved in vesicle trafficking.

### CRYSP and CRONE are crystalloid proteins essential for sporozoite development

Western blot analyses using an anti-HA antibody detected a ∼ 33 kDa CRYSP::3XHA protein in extracts from purified *in vitro* cultured *crysp::3xha* ookinetes and, to a much lower levels, from gametocytes, both prior to and after induction of gametogenesis **(Figure 4A)**. Similarly, the ∼ 33 kDa CRONE::3×HA protein was detected in ookinete and, less so, gametocyte extracts of the *crone::3xha* line **(Figure 4B)**.

**Figure 4.**
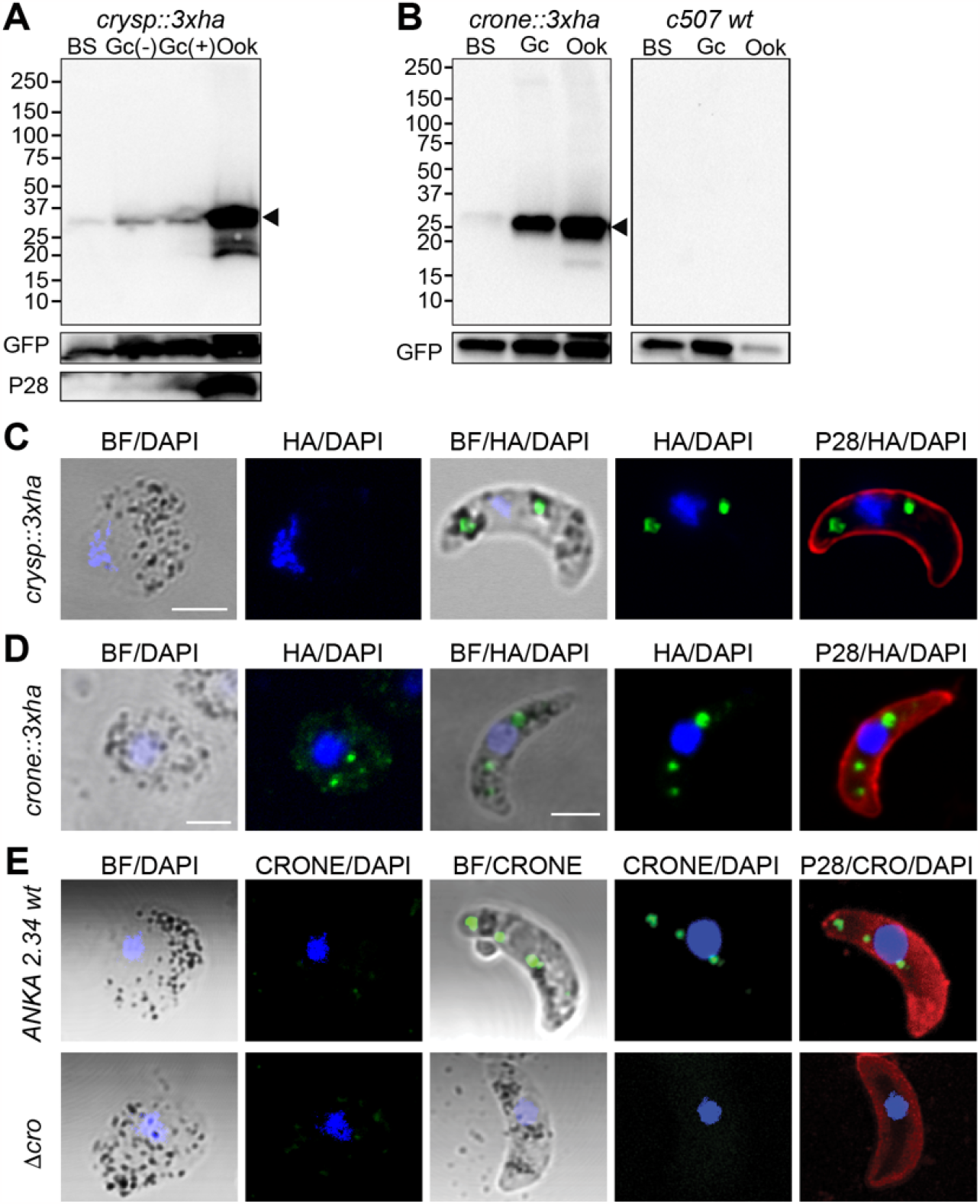
CRYSP and CRONE protein expression and localization. **(A)** Western blot analysis of whole cell lysates of the *crysp::3xha* parasite line using an α-HA antibody under reducing conditions. The CRYSP::3XHA specific band is indicated with a black arrowhead. GFP and P28 were used as loading and stage-specific controls, respectively. BS, mixed blood stages; Gc(-), inactivated and purified *in vitro* cultured gametocytes; Gc(+), activated and purified *in vitro* cultured gametocytes; Ook, purified *in vitro* cultured ookinetes. **(B)** Western blot analysis of whole cell lysates of the *crone::3xha* (left) and *c507 wt* control (right) parasite lines using an α-HA antibody under reducing conditions. The CRONE::3XHA specific band is indicated with a black arrowhead. GFP and P28 were used as loading and stage-specific controls, respectively. BS, mixed blood stages; Gc, purified *in vitro* cultured gametocytes; Ook, purified *in vitro* cultured ookinetes. **(C)** Immunofluorescence assays of *crysp::3xha* purified gametocytes (Gc) and ookinetes (Ook) of the *crysp::3xha* parasite line, stained with α-HA (green) and α-P28 (red) antibodies. DNA was stained with DAPI (blue). BF; brightfield. Scale bar is 2.5 μM. **(D)** Immunofluorescence assays of *crone::3xha* purified gametocytes (Gc) and ookinetes (Ook), stained with α-HA (green) and α-P28 (red) antibodies. DNA was stained with DAPI (blue). BF; brightfield. Scale bar is 2.5 μM. **(E)** Immunofluorescence assays of *2*.*34 wt* purified gametocytes and ookinetes, stained with α-CRONE (green) and α-P28 (red) antibodies. *Δcro* parasites were used as negative controls. DNA was stained with DAPI (blue). BF; brightfield. Scale bar is 2.5 μM.

We examined the cellular localization of the two proteins in immunofluorescence assays of gametocytes and ookinetes **(Figure 4C-D)**. In both cases, a clear and distinct spot pattern that always colocalized with the hemozoin (visible in bright field) was detected in the ookinete. This pattern is the hallmark of crystalloid localization in *P. berghei* (Carter et al., 2008). Multiple ookinete observations revealed that the number of spots varied from 1 to 3, which were always in association with the hemozoin containing vesicles. The two proteins were henceforth named CRONE for “crystalloid oocyst not evolving” and CRYSP for “crystalloid needed for sporozoites”.

In the *crone::3xha* line, a vesicle-like albeit less prominent staining pattern was also detected in the female gametocytes, consistent with the high CRONE protein abundance in gametocyte extracts. Crystalloids are organelles known to be specific to ookinetes and young oocysts, thought to form soon after fertilization through active assembly of endoplasmic reticulum (ER)-derived vesicles (Dessens et al., 2021). Some of the known crystalloid proteins are also synthesized in the gametocytes (Saeed et al., 2015). Therefore, one can speculate that the CRONE::3×HA-stained gametocyte vesicles are crystalloid precursor subunits. Whilst this may be true, the expression of CRONE in gametocytes could be due to the CRONE::3×HA expression design that used the *P. berghei* dihydrofolate reductase (DHFR) 3′ untranslated region (UTR). Cis-acting elements in the 5′ UTR or 3′ UTR of DOZI-regulated genes have been shown to be important for translational repression (Braks et al., 2008). Indeed, a previous study that expressed a GFP-tagged version of CRONE using the 3′ UTR of P28 that is also translationally repressed by DOZI found that GFP is restricted to the ookinete crystalloid (Guerreiro et al., 2014). To examine this, we raised rabbit polyclonal antibodies against a codon-optimized fragment of CRONE (amino acids 24-235) expressed in *Escherichia coli* cells. Using these antibodies in immunofluorescence assays, we detected a clear ookinete crystalloid signal, but this signal was absent from gametocytes **(Figure 4E)**. This indicated that the gametocyte signal detected in the *crone::3xha* line is likely due to leaky DOZI post-transcriptional repression.

Our findings add to the increasing recognition of the pivotal role of the crystalloid in sporogony and mosquito-to-human transmission. Although the details remain poorly understood, the current view is that the crystalloid assembles from ER-derived vesicles in a microtubule-dependent mechanism and serves in transporting functionally diverse proteins to the maturing oocyst (Dessens et al., 2021; Saeed et al., 2015). The commonly more than one and often two ookinete crystalloids appear as a single organelle in the oocyst (Saeed et al., 2015), but it is unclear whether this is already a single multi-lobed organelle or is due to asynchronous dissolution or merging of separate crystalloids. The reason behind the transportation of proteins by this organelle instead of their contemporaneous synthesis in the oocyst is unclear; it may be attractive to speculate that during early stages of development, the metabolic environment in the oocyst is incompatible with *de novo* transcription and translation of proteins needed for sporogenesis. An alternative hypothesis is that this organelle functions in the ookinete and young oocyst with a knock-on effect in the mature oocyst.

The crystalloid founding molecules are the LCCL-lectin adhesive-like proteins (LAPs) that exhibit modular domain architectures implicated in protein, lipid and/or carbohydrate binding (Dessens et al., 2021). A recent proteomic analysis of *P. berghei* crystalloids revealed that the LAPs are part of an extended protein interaction network, which includes CRYSP and TPM2 (PBANKA_1104100), a structural homologue of CRONE (Tremp et al., 2020). Both CRONE and TPM2 contain a TPM domain, named after its founding proteins, the *Arabidopsis thaliana* TLP18.3 and Psb32 and the *Caenorhabditis elegans* MOLO-1, as well as a C-terminal transmembrane domain. The TPM domain, despite being structurally conserved, is made of diverse amino acid sequences indicating varied functions. Indeed, the conserved amino acids required for the phosphatase activity of TLP18.3 (Wu et al., 2011) are absent from CRONE and TPM2 suggesting different functions of these crystalloid proteins. It is yet unclear whether TPM2 shares the same phenotype as CRONE, *i*.*e*., normal mitotic replication but failure of sporulation.

The discovery of CRYSP brings a new perspective into the role of the crystalloid, as this is the first crystalloid protein unambiguously shown to be involved in sporozoite egress from the oocyst or infectivity rather than formation. A similar function has been previously suggested for the *P. falciparum* LAP orthologs, CCp2 and CCp3, but that study has not examined whether the seemingly normal oocyst sporozoites as seen with electron microscopy are fully formed and can be separated from the body of the oocyst (Pradel et al., 2004). Indeed, this study did not detect sporozoites in the mosquito hemocoel. Disruption of LAP4, the *P. berghei* ortholog of PfCCp2, exhibits abnormal crystalloid biogenesis and gives rise to small and early sporulating oocysts that produce non-infectious sporozoites (Saeed et al., 2018). In contrast, disruptions of LAP1, the *P. berghei* ortholog of PfCCp3, and of LAP3 produce parasites that are devoid of crystalloids and fail to complete oocyst maturation (Carter et al., 2008; Tremp et al., 2017).

In addition to the LAPs, CRONE and CRYSP, the other two characterized crystalloid *P. berghei* proteins are the palmitoyl-S-acyl transferase, DHHC10, thought to be involved in post-translational lipid modification of proteins (Santos et al., 2016), and the NAD(P) transhydrogenase, NTH, that generates NADPH (Tremp et al., 2020). Both proteins are predicted to be transmembrane and required for crystalloid biogenesis and sporozoite formation. Therefore, a common theme that emerges from these studies is that biogenesis of the crystalloids is dependent on the successful loading of most if not all of their protein cargo. This would suggest that the sporogony-deficient phenotype of the aforementioned mutants is an all-encompassing effect linked to the lack of the crystalloids rather than each individual protein. It remains to be seen whether this is true for CRONE and CRYSP, although the distinct phenotype of the latter suggests otherwise.

## Methods

### Resources and reagents sharing

All requests for resources and reagents may be directed to Dina Vlachou. In laboratory stocks and reagents, the various genes studies here are often referenced with their temporary given codes, i.e., *PBANKA_0413500* (*STONES*) is N350, *PBANKA_1338100* (*CRYSP*) is N38, *PBANKA_1353800* (*ROVER*) is c53, and *PBANKA_0720900* (*CRONE*) is c72.

### Ethics statement

Animal procedures were reviewed and approved by the Imperial College Animal Welfare and Ethical Review Body (AWERB) and carried out in accordance with the Animal Scientifics Procedures Act 1986 under a UK Home Office project license.

### Parasite lines, cultivation, and mosquito infections

*P. berghei* lines used were: the wildtype cl15cy1 line (ANKA 2.34); the constitutively GFP-expressing and selectable marker free 507m6cl1 (c507) line, which has the GFP under the control of the ef1a promoter and is integrated into the 230p (PBANKA_0306000) gene locus (Janse et al., 2006); the non-gametocyte producer ANKA 2.33 line (Sinden et al., 1996); a selectable marker free HAP2 knockout line generated using the PbGEM-102303 vector and the parental reference line *1804cl1* (RMgm928) which expresses mCherry under the control of the HSP70 promoter (Burda et al., 2015); and the Nek4 knockout line *826cl1* (RMgm257). The cultivation and purification of parasites were carried out as described (Sinden et al., 1996). Feeding of *A. coluzzii* mosquitoes was done by direct feeding on mice with parasitemia of 5-6% and gametocytemia of 1-2% or by feeding on *in vitro* cultured ookinetes.

### DNA microarray hybridizations

The *P*.□*berghei* Agilent oligonucleotide microarray platform has been described previously (Akinosoglou et al., 2015). Remapping of oligonucleotide probes on the latest *P*. □ *berghei* genome assembly and annotation release of PlasmoDB (version 35, released on 09/2022). 30–40 *A*. □ *coluzzii* midguts from 3 biological replicate infections with ANKA 2.34 and ANKA 2.33 were dissected at 1 □h pbf in ice cold phosphate-buffered saline (PBS) and immediately immersed in TRIzol reagent (Invitrogen). Total RNA was extracted according to the manufacturer’s instructions and quantified using NanoDrop^®^ ND-1000 Spectrophotometer (Thermo Scientific). 2 □μg of total RNA were used for the generation and labelling of cRNA using the Agilent low RNA input fluorescence amplification kit according to manufacturer’s instructions. 2 □μg of Cy3 (ANKA 2.33) and Cy5 (ANKA 2.34) labelled cRNA were mixed and hybridized on the microarrays using the Agilent *in situ* hybridization kit according to the manufacturer’s instructions. After washing, the hybridized microarrays were scanned using the Gene-Pix 4000B scanner (Axon Instruments). Grid alignment, registering spot signal intensity, estimation of local backgrounds and manual inspection of spot quality were carried out using Gene-Pix Pro 6.1. Data normalization was carried out using the locally weighted linear regression method (Lowess) in GeneSpring GX 12.6 (Agilent Technologies). Significant transcriptional differences stages were calculated using a one-way ANOVA with a *P*-value cut-off of 0.05, following correction with the Benjamini-Hochberg hypergeometric test.

### Generation of pools of transgenic parasites

To generate pools of transgenic parasites, *Plasmo*GEM vectors were combined in equal concentrations, and the mixture was digested with NotI to linearize the targeting vector. A total of 3.2 μg of the purified digested vector mix containing about 100 ng of each vector was used per transfection as previously described (Gomes et al., 2015). Briefly, purified schizonts derived from the *hap2ko* mCherry parasite was electroporated using the FI-115 program of the 4D nucleofector system (Lonza). Transfected schizonts were then injected intravenously into BALB/c mice, and drug selection of resistant parasites was carried out by the administration of pyrimethamine in the drinking water (70 μg/mL). Mouse infections following transfection was monitored daily by Giemsa staining of tail blood smears.

### Mosquito transmission of transgenic parasite pools

At 8 days post transfection with the *Plasmo*GEM vector pool when mice parasitemia was 6-8%, blood was obtained from infected mice via heart puncture and mixed with blood derived from mice infected with the NEK4 knockout parasite with the same parasitemia in a ratio of 2:1. This mixture was used to infect BALB/c mice. At a parasitemia of 3-4%, mice were used in direct mosquito feeds. 30-50 mosquito midguts and salivary glands per biological replicate were dissected and collected for genomic DNA extraction.

### Library generation and barcode sequencing

Genomic DNA was extracted from mouse blood sampled at day 4-8 post infection with the transfected parasites and at day 4 post infection with serially passaged parasites, and from mosquito midguts and salivary glands at days 10-12 and 21 pbf, respectively, using phenol-chloroform extraction. *Plasmo*GEM vector specific barcodes were sequenced using Illumina MiSeq as described previously (Gomes et al., 2015). Briefly, *Plasmo*GEM barcodes were amplified by PCR using the genomic DNA and the primers arg444 and arg445 (**Table S8**). The PCR amplicons were then used for a second PCR that introduced 5′ adaptors and multiplexing barcodes using primers shown in **Table S8**, and the resulting libraries were pooled at 100 ng per library and sequenced using the Illumina MiSeq Reagent Kit v2. After sequencing, barcode sequences were extracted from the output raw sequence file using a Perl script, counted, and their relative abundance (counts per 1000 barcodes) within each pool was determined. Statistical analysis was performed using a student’s t-test followed my multiple testing correction.

### Generation of single knockout transgenic parasites

For the generation of the *Δmap2* and *Δhap2* background lines, we used the *Plasmo*GEM vectors PbGEM-111778 and PbGEM-102303, respectively. A total of 5 μg of each plasmid was used to transfect segmented *P. berghei* schizonts as previously described (Fang et al., 2018). Briefly, schizonts were electroporated using the FI-115 program of the Amaxa Nucleofector 4D, after which parasites were immediately injected intravenously into the tail vein of BALB/c mice. Transgenic parasites were selected with 0.07 □mg/mL pyrimethamine (Sigma) in drinking water from day 1 pi. Disruption was confirmed in the resistant parasite populations by PCR and clonal lines were derived by limiting dilution. To allow the use of *Δmap2* and *Δhap2* as background lines in the screen, we induced excision of the resistance cassette from the genome using negative selection, through the administration of 5 fluorocytosine (1□mg/mL, Sigma) via the drinking water (Braks et al., 2006). This was possible because each resistance cassette carried a gene encoding the yFCU that counteracts the administered 5 fluorocytosine. Each mutant was finally re-genotyped to confirm correct excision of the resistance cassette and clonal lines were once again derived by limiting dilution.

For disruption of *STONES* and *CRYSP*, we used the PbGEM_230494 and PbGEM_058356 *Plasmo*GEM vectors, respectively. The targeting cassettes were released by NotI digestion resulting in 84% and 80% deletion of the CDS of *STONES* and *CRYSP* at the 5′ end. Partial (66%) knockout of *CRONE* and full knockout of *ROVER* and *SPM1* was carried out by double crossover homologous recombination in the c507 line. For this, EcoRI/BamHI 5′ homology arms and Apa/HindIII 5′ homology arms were amplified from genomic DNA using the primer pairs P1/P2 (588 bp), P5/P6 (728 bp) and P9/P10 (620 bp) and P3/P4 (573 bp), P7/P8 (558 bp) and P11/P12 (648 bp), respectively (**Table S9**). These fragments were cloned into the Pbs-TgDHFR vector with homology arms flanking a modified *Toxoplasma gondii* dihydrofolate gene (*TgDHFR/TS)* cassette that confers resistance to pyrimethamine. The gene targeting cassettes were released by ApaI/BamHI digestion. Transfection, drug selection of transgenic parasites and clonal selection by dilution cloning was carried out as previously described (Janse et al., 2006).

### Generation of tagged transgenic parasites

For the C-terminal 3xHA tagging of *STONES, CRYSP* and *CRONE* in the *c507* line, we used the *Plasmo*GEM vectors PbGEM012712, PbGEM058364 and PbGEM089977, respectively (Gomes et al., 2015). The C-terminal 3×HA tagging of *ROVER* in the c507 line was generated by Gibson assembly. Firstly, a 694 bp 5′ homology arm ApaI fragment corresponding to the most 3′ region of the CDS and the 3XHA sequence was amplified using the primer pairs P30/P31 (**Table S9**). The 460bp *DHFR 3*′*UTR* SacII fragment was amplified from the pL00018 vector (MRA-787, MR4) using the primers P32/P33. An overlap PCR using both fragments was set up to generate the Apa/SacII *ROVER::3XHA::DHFR 3*′*UTR*. A 560 bp Xho/XmaI 3′ homology arm region corresponding to the *3*′*UTR* of the gene was amplified using the primer pairs P34/P35 (**Table S9**). The *ROVER* fragments were cloned flanking the *hDHFR/yFCU* selection cassette into plasmid pL0035 (Braks et al., 2006).

### Genotypic analysis of transgenic parasites

Following drug selection and clonal selection, parasite genomic DNA was extracted from blood sampled from parasite positive mice using the DNeasy kit (Qiagen). Successful gene modification events or maintenance of the wildtype locus was performed by PCR using primers listed in Table S9.

### Genetic crosses

Genetic crosses between the *Δrov* or *Δc57* and the *Δhap2* (male-deficient) or *Δnek4* (female-deficient) lines were carried out by infecting mice with different combinations of mutant parasites, which were then used for direct feeding of *A*. □ *coluzzii* mosquitoes.

### Exflagellation assays

Blood from infected mice exhibiting 1-2% gametocytemia was added to RPMI medium (RPMI 1640, 20% v/v FBS, 100 μM xanthurenic acid, pH 7.4) in a 1:40 ratio and incubated for 10 min. Exflagellation events were counted in a standard hemocytometer under a light microscope.

### Macrogamete to ookinete conversion assays

Ookinete formation was assessed by conversion assays. Blood from infected mice exhibiting 1-2% gametocytemia was added to RPMI medium (RPMI 1640, 20% v/v FBS, 100 μM xanthurenic acid, pH 7.4) and incubated for 24 hours at 21°C to allow for ookinete formation. This suspension was then incubated with a Cy3-labelled 13.1 mouse monoclonal α-P28 (1:50 dilution) for 20 min on ice. The conversion rate was calculated as the percentage of Cy3 positive ookinetes to Cy3 positive macrogametes and ookinetes.

### Ookinete motility assays

A 24-hour *in vitro* culture of mature ookinetes was added to Matrigel (BD biosciences), and the mixture was dropped onto a slide and allowed to set at room temperature for 30 min. Time-lapse microscopy (1 frame every 5 seconds, for 10 min) of ookinetes were taken on a Leica DMR fluorescence microscope and a Zeiss Axiocam HRc camera controlled by the Axiovision (Zeiss) software. The speed of individual ookinetes was measured using the manual tracking plugin in the Icy software package.

### Invasion assay

Total RNA was extracted from *A. coluzzii* midguts infected with *P. berghei* 24 hours pbf using the TRIzol reagent (Invitrogen). The RNA was used to generate cDNA that was subsequently used in the amplification of *CTL4* using primers P51/P52 with T7 overhangs to produce double-stranded RNA using the T7 high yield transcription kit (ThermoFisher). The double-stranded RNA was purified using the RNeasy kit (Qiagen) and 0.2 μg in 69 nL was injected through the mesothoracic spiracle into the hemocoel cavity of *A. coluzzii* mosquitoes using glass capillary needles and the Nanoject II microinjector (Drummond Scientific). Injected mosquitoes were allowed to recover for 3 days before blood feeding.

### *P. berghei* mosquito infections

Mosquitoes were infected by direct feeding on mice infected with *P. berghei* at a parasitemia and gametocytemia of 5-6% and 1-2%, respectively. To determine oocyst load, midguts were dissected at 7-10 days pbf and fixed in 4% paraformaldehyde. Melanized parasites and oocyst numbers were counted under a light and fluorescent microscope, respectively. To determine sporozoite load, 25-30 midguts and salivary glands were dissected 15 and 21 days pbf, respectively, homogenized and sporozoites counted in a standard hemocytometer under a light microscope. To assess mosquito-to-mouse transmission, about 30 *A. coluzzii* mosquitoes that had blood-fed on *P. berghei*-infected mice 20-22 days earlier were allowed to feed on 2-3 anaesthetized C57/BL6 mice. Mouse parasitemia was monitored until 14 days post mosquito bite by Giemsa staining.

### Ookinete injection

The concentration of ookinetes obtained from a 24-hour *in vitro* ookinete culture was adjusted to achieve injection of approximately 800 ookinetes per mosquito delivered through injection of *A. coluzzii* females through the mesothoracic spiracle using glass capillary needles and the Nanoject II microinjector (Drummond Scientific). Salivary glands were dissected 21 days post injection and homogenized, and sporozoite numbers were counted using a standard hemocytometer under a light microscope.

### CRONE expression, purification and antibody production

A *CRONE* CDS fragment corresponding to amino acids 23-235, which excludes the predicted N-terminal signal peptide and C-terminal transmembrane domain, was codon optimized for expression in *E. coli* (GeneArt, ThermoFisher). This fragment was amplified with the primer pair P49/P50 (**Table S9**) and cloned into a NotI digested pET-32b protein expression vector, which carries the N and C-terminal 6xHistidine tags (Novagen), using the Hi-Fi DNA assembly kit (New England Biosciences).

*E. coli* BL21 cells (New England Biosciences) containing the recombinant protein expression plasmid were grown at 37°C and induced with 1 mM isopropyl-1-thio-β-d-galactopyranoside at 18°C for 16 hours. Cells were harvested and lysed using cell lytic (Sigma) containing the protease inhibitors cOmplete EDTA-free (Roche). Cell debris were removed by centrifugation. The His-tagged CRONE protein was purified by cobalt affinity chromatography using TALON® metal affinity resin (Takara) under native conditions in phosphate buffered saline (PBS), pH 7.4. Protein samples were analyzed by SDS-PAGE to determine purity prior to their use for immunization in rabbits for the generation of an affinity purified polyclonal antibody (Eurogentec).

### Western blot analysis

Western blot analysis was performed on whole cell lysates and fractionated cell samples. To extract whole cell lysates, purified blood stages, gametocytes and ookinetes were suspended in whole cell lysis buffer (1XPBS, 1% v/v Triton X-100). For fractionation, firstly, the soluble fraction was obtained by suspension and homogenization of purified parasites in soluble cell lysis buffer (50mM Tris, 300mM NaCl). Secondly, to obtain the Triton-Soluble fraction, the pellet from the prior treatment was then resuspended and homogenized in Triton-solubilization buffer (50mM Tris, 300mM NaCl, 1% v/v Triton X-100). Finally, to obtain the Triton-Insoluble fraction, the pellet from the prior treatment was resuspended and homogenized in Laemilli SDS buffer. Protein fractions were boiled under reducing conditions and separated using 4-20% sodium dodecyl sulfate (SDS) polyacrylamide gel electrophoresis. The gel separated proteins were transferred to a polyvinylidene difluoride (PVDF) membrane. Proteins were detected using rabbit α-HA (Cell Signaling Technology) (1:1000), goat α-GFP (Rockland chemicals) (1:1000) and 13.1 mouse monoclonal α-P28 (1:1000) antibodies. Secondary horseradish peroxidase (HRP) conjugated goat α-rabbit IgG, goat α-mouse IgG antibodies (Promega) and donkey α-goat IgG (Abcam) were used at 1: 2,500, 1: 2,500 and 1: 5,000 dilutions, respectively. All primary and secondary antibodies were diluted in 5% w/v milk-PBS-Tween (0.05% v/v) blocking buffer.

### Indirect immunofluorescence assays

Blood stage gametocytes, ookinetes and sporozoites were fixed in 4% paraformaldehyde (PFA) in PBS for 10 min at room temperature. Fixed parasites were washed 3X with 1XPBS for 10 min each and then smeared on glass slides. Permeabilization of the parasites was done using 0.2% v/v Triton X-100 in PBS for 10 min at room temperature. Permeabilized parasites were washed 3 times in PBS for 10 min each and then blocked with 1% w/v bovine serum albumin in PBS for 1 hour at room temperature. Parasites were stained with rabbit α-HA (CST) (1:1000), rabbit α-GFP (ThermoFisher) (1:500), 13.1 mouse monoclonal α-P28 (Ref) (1:1000) and 2A10 mouse monoclonal α-PfCSP (1:100) antibodies. Alexa Fluor rabbit 488 and mouse 568 conjugated secondary goat antibodies (ThermoFisher) were used at a dilution of 1:1000. 4′,6-diamidino-2-phenylindole (DAPI) was used to stain nuclear DNA. Images were acquired using a Leica SP5 MP confocal laser-scanning microscope. Images were visualized using Image J.

### Statistical analysis

All statistical analysis was performed using GraphPad Prism v8.0. P-values for exflagellation, ookinete conversion and motility assays were calculated using a two-tailed, unpaired Student’s *t*-test. For statistical analyses of the oocyst or melanized parasite infection intensities and presence of oocysts (infection prevalence), *P-* values were calculated using the Mann-Whitney test.

## Acknowledgements

We thank Claudia Wyer and Temesgen Menberu Kebede for assistance with mosquito and parasite culturing and Nikolaos Trasanidis for assistance with the transcriptome analysis. We also thank Luc Duchateau for suggestions on statistical analyses of the STM screen data. The work was funded by a Wellcome Trust Investigator Award to GKC (107983/Z/15/Z) and a Medical Research Council (MRC) project grant (MR/T000929/1) to GKC and DV. AB was supported by a Wellcome Trust PhD fellowship award to Imperial College London (102126/B/13/Z).

## Author contributions

Conceptualization: OB, DV and GKC; Methodology: CVU, ARG, OB, DV and GKC; Validation: CVU, DV and GKC; Formal analysis: CVU, ARG, DV and GKC; Investigation: CVU, ARG, MG, MC, AB, TRBB and DV; Resources: OB, DV and GKC; Data Curation: CVU, DV and GKC; Writing - Original Draft: CVU, ARG, DV and GKC; Writing - Review & Editing: DV and GKC; Visualization: DV and GKC; Supervision: DV and GKC; Project administration: DV and GKC; Funding acquisition: DV and GKC.

