## Supplemental materials for "Reverse genetic screen identifies malaria parasite genes required for gametocyte-to-sporozoite development in its mosquito host"

#### Supplementary Information

##### *P. berghei* gametocyte transcriptome

Labelled RNA from three independent replicate infections of *A. coluzzii* with either the *P. berghei* ANKA 2.34 or the *P. berghei* ANKA 2.33 was used in competitive hybridizations on a 4X44K *P. berghei* DNA microarray (Akinosoglou et al., 2015). Oligonucleotide remapping showed that the microarray encompassed 4,288 of the 5,254 genes predicted in the *P. berghei* genome according to PlasmoDB version 35, released on 09/2022. Of the 4,288 genes, 3,428 genes were the same as in the original array design, thus were represented by the same probes. The remaining 860 genes were because of merging or splitting of genes between various gene builds and were therefore represented by a different combination of probes compared with the original probe combinations. They were not considered further to prevent errors due to gene annotation problems. Therefore, the final gene coverage obtained was 65%.

Data normalization and one sample t-test on gene expression data showed that 274 genes exhibited statistically significant regulation with at least 1.6-fold difference in the *P. berghei* *wt* infected midguts compared to the *NGP* infected midguts (**Table S1**). The analysis showed that 69% (189 genes) and 31% (85 genes) are up and downregulated respectively. In the downregulated genes, a good correlation is observed between the presence of transcript in gametocytes and the presence of transcripts in prior ABS’s. 87% of these genes are expressed in ABS’s (Hall et al., 2005; Le Roch et al., 2003; Otto et al., 2014). On the other hand, only 26% of the upregulated genes have transcripts or proteins reported to be expressed in ABS’s.

Domain analysis of the upregulated genes showed that 8% (16 genes) encode proteins associated with cell signalling functions and includes the zygote developmental genes NEK2 (Reininger et al., 2005) and protein kinase 7 (PK7) (Tewari et al., 2010). 17% of genes encode proteins with cytoskeletal functions and 10 of these genes encode for pellicular proteins and include the inner membrane complex proteins. 13 genes encode proteins that function in either DNA replication, mitosis, meiosis or DNA repair. Examples include the nuclear formin-like male inherited sporulation factor important for transmission (MISFIT) (Bushell et al., 2009) and the meiotic recombination protein DMC1 (Mlambo et al., 2012). Proteins with a GINS domain or LISH domain or structural maintenance of chromosomes (SMC) domain also belong to this group. 10 genes encode proteins associated with gene regulation either at the transcriptional or post-transcriptional level and include DOZI and various zinc finger containing proteins. The fifth group includes 24 genes with defined domains that do not fit into groups 1-4 but have been however implicated in sexual and sporogonic development. MDV1, PPLP2, GEST and PF16 function in gametogenesis (Deligianni et al., 2013; Ponzi et al., 2009; Straschil et al., 2010; Talman et al., 2011; Wirth et al., 2014). P25, P28 and PPLP3 are involved in ookinete midgut invasion (Ishino et al., 2006; Tomas et al., 2001). A group of CPW-WPC domain containing proteins were also put into this group. Most of the proteins belonging to this group have domains associated with adhesive properties. 16% of genes are associated with general processes such as metabolism and includes various enzymes and transporters. A group of 4 genes do not fit into groups 1-6 and appear to be mainly ABS genes. Almost 31% of genes (60 genes) do not have yet, known domains and are annotated as genes with unknown functions.

Transcript and/or proteins have been detected in previous studies (Akinosoglou et al., 2015; Florens et al., 2002; Hall et al., 2005; Howick et al., 2019; Lasonder et al., 2008; Otto et al., 2014) at the ookinete and/or sporogonic developmental for roughly half (45%) of the upregulated genes. Finally, 60% (115 genes) of the upregulated genes are downregulated in the absence of translational repression mediators DOZI and CITH (Mair et al., 2006; Mair et al., 2010). It is important to note that due to the microarray coverage, about 1800 genes were not investigated. RNA-seq data (Otto et al., 2014) comparing schizont (ABS) and gametocyte expression, showed that 380-450 of these genes were at least 2-fold upregulated in the gametocyte in one (452) or two (380) experiments. Investigation for gametocyte DOZI/CITH transcript regulation which can be used as a putative marker for protein requirement in the mosquito stage (Guerreiro et al., 2014) showed that only 5% of these transcripts were DOZI/CITH regulated (Mair et al., 2006; Mair et al., 2010). However, this does not exclude the remainder genes, as lack of DOZI regulation does not necessarily equate to non-necessity of these genes for later stages of development in the mosquito (Carter et al., 2008; Lin et al., 2013; Raine et al., 2007). Transcript abundance of these gametocyte genes in the ookinete may point to a function in later mosquito stage development. 38 genes have fragments per kilobase of transcript per million mapped reads (FPKM) above 300, 59 genes have FPKM between 100-300 and 355 genes have FPKM less than 100 (Otto et al., 2014).

##### Genes with known phenotypes included in the screen

Included in the screen were genes with known phenotypes included GAMER and HADO that are important for ookinete formation (Akinosoglou et al., 2015), SOAP that encodes a secreted ookinete adhesive protein involved in midgut invasion (Dessens et al., 2003), PIMMS43 that is essential for the ookinete-to-oocyst transition and protects ookinetes from a mosquito systemic immune response (Ukegbu et al., 2020), PIMMS2 (aka SUBO) that is essential for ookinete midgut egress (Ukegbu et al., 2017), PIMMS01 and PIMMS57 involved in ookinete-to-oocyst transition (Ukegbu et al., 2021), PBANKA_0720900 (named here CRONE) that encodes a crystalloid-related protein essential for oocyst development (Guerreiro et al., 2014), TRAP that encodes a sporozoite micronemal protein (Spaccapelo et al., 1997), and the ookinete and sporozoite specific G2 protein (Tremp et al., 2013). Additionally, there were nine genes encoding members of the 6-cys family (van Dijk et al., 2005; van Dijk et al., 2001; van Dijk et al., 2010), of which some are already known to be associated with either the gametocyte-to-ookinete developmental transition or with *Plasmodium* developmental progress in the mammalian host liver: P47, P48/45, P230, P230p, P38, P12p, B9, P36 and P36p. PV1 (Chu et al., 2011), a gene exclusively expressed in ABS and shown to be associated with the *P. falciparum* intraerythrocytic growth was included as a control. Vectors for epitope tagging of four genes (PBANKA_0306000, PBANKA_0410400, PBANKA_0514900 and PBANKA_1002400) previously shown not to be detrimental for parasite growth were included as controls.

#### Supplementary figures

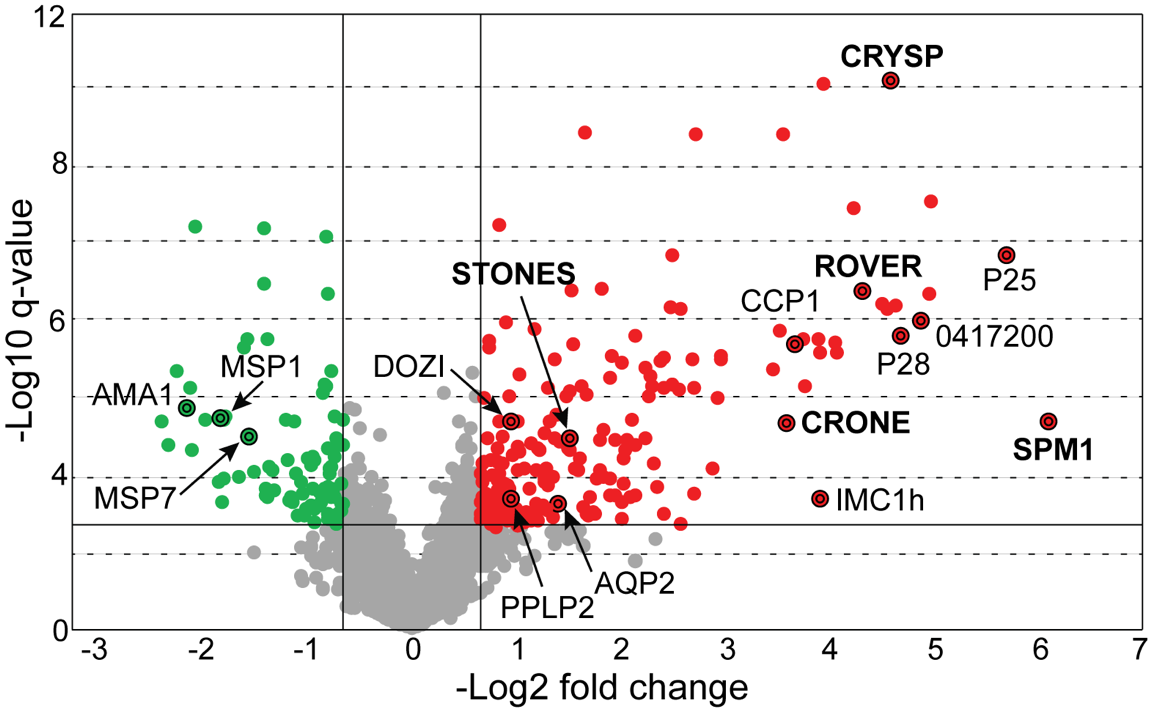

##### Figure S1. Transcriptional enrichment in *P. berghei* gametocytes.

Volcano plot showing differential gene expression measurements between *wt* and the *NGP* *P. berghei* line 1 hpi in *A. coluzzii* midgut. Red and green dots represent the 274 genes that are regulated by at least 1.6-fold, respectively, in the *wt* line compared to the *NGP* line. Grey dots represent genes that do not show significant regulation. 189 genes (red dots) were enriched in the wild type compared to the non-gametocyte producing parasite line and include genes involved in sexual and sporogonic development such as *PPLP2, DOZI, P25,* P28, *IMC1h* and *LAP2*. Eighty-five genes (green) were identified to be downregulated, and include genes encoding putative blood stage proteins such as AMA1 and the merozoite surface proteins MSP’s 1and 7. The genes in bold represent genes selected for further characterization.

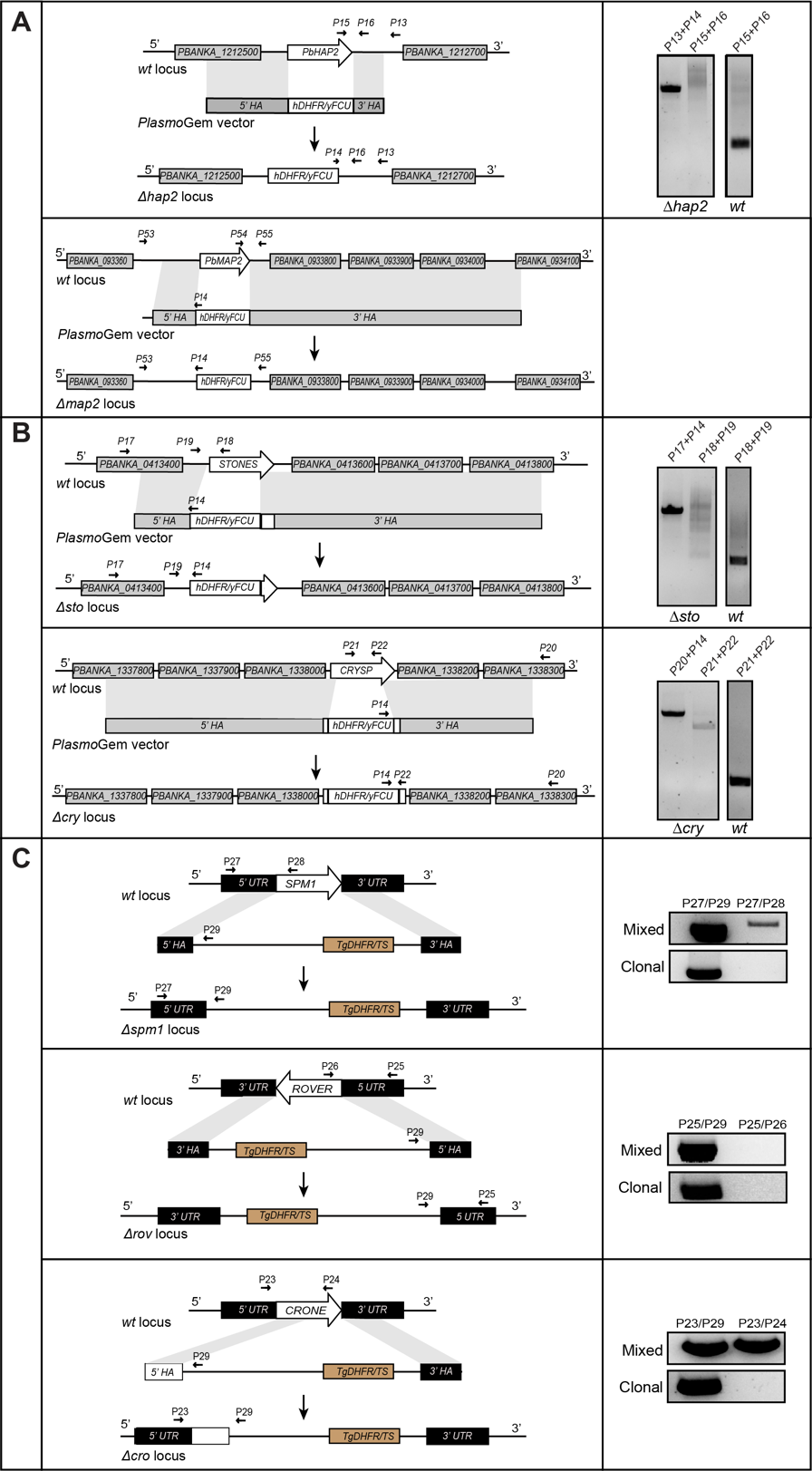

##### Figure S2. Generation of mutant parasites

Schematic representation of the disruption of (**A**) *HAP2* and *MAP2* in the *1804cl1* line and (**B**) *STONES* and *CRYSP* in the *c507* line, using *Plasmo*GEM vectors carrying the *human DHFR* (*hDHFR)/yFCU* pyrimethamine resistance cassette, as well as of **(C)** *ROVER*, *CRONE* and *SPM1* in the *c507* line by using vectors that carry a modified *Toxoplasma gondii DHFR (TgDHFR)* pyrimethamine resistance expression cassette in the *c507* line**.** In each panel, the *wt* genomic locus (top), the transfected DNA fragment carrying the homologous arms (HA) and the selectable markers (middle), and the final transgenic locus after gene disruption by double crossover homologous recombination (bottom) are shown. Constructs are not drawn to scale. Small arrows show the primers used for the confirmation of integration of gene targeting constructs and deletion of the gene of interest using PCR as shown in the panels on the right. The *1804cl1* and *c507 wt* parental parasites were used as a control for detection of the *wt* locus.

```
PbSTONES      MKKEEEFLKNEIEKESIALFKKNLLKESYKWFKDTDYFVKIKNDNNYNNIENKTETKNIV 60
PySTONES      MKKGEEFLKNEIEKESIALFKKNLLKESYKWFKDTDYFVKIKNDNNYNNIENKTETKNIV 60
PcSTONES      MKKEEEFLKNEIEKESIALFKKNLLKESYKWFKDTDYFVKIKNDNNYNNIENKTETKNIV 60
PfSTONES      MRKDESEMKKKIEKESIALFKKNLLKESYKWFKDTDYFVKLKNENKYSNIENKTETRNIV 60
PvSTONES      MRARDPSLKSQIEKESIALFKKNLLKESYSWFKDTDYFEKLKSEKKYSNAENERETKNVV 60
PkSTONES      MKARDPSLKSQIEKESIALFKKNLLKESYNWFKDTDYFEKLKSENKYSNGENEHETKNVV 60
Consensus     *:  :  :*.:******************.******** *:*.:::*.* **: **:*:*

PbSTONES      TGRVQLGWGGTFFLLCCVVVICDNSGWQSCNHYNNNIFYSPESLTIVYSFLYVIYAFPLF 120
PySTONES      TGRVQLGWGGTFFLLCCVVVICDNSGWQSCNHYNNNIFYSPESLTLVYSFLYAIYAFPLF 120
PcSTONES      TGRVQLGWGGTFFLLCCVVVICDNSGWQSCNHYNNNIFYSPESLTIVYSFLYVIYAFPLF 120
PfSTONES      TGRLQLGWGGTFFLLCCVVVICDNSGWQSCNHYNNNIFYSPESLTIVYSFLYIIYGVPLF 120
PvSTONES      TGRVQLGWGGTFFLLCCVVVICDNSGWQSCNHYNNSIFYSPEALTVVYSILFIVYAFPLF 120
PkSTONES      TGRVQLGWGGTFFLLCCVVVICDNSGWQSCNHYNNSIFYSPEALTVVYSILFIVYAFPLF 120
Consensus     ***:*******************************.******:**:***:*: :*..***

PbSTONES      IFHQCLGLVSQGNYVKSCRMLSPHMGILTIFSLFGTITFCAKEICNFSAEFLQNIIRLYN 180
PySTONES      IFHQCLGLVSQGNYVKSCRMLSPHMGILTIFSLFGTITFCAKEICNFSAELLQNIIRLYN 180
PcSTONES      IFHQCLGLVSQGNYVKSCRMLSPHMGILTIFSLFGTITFCAKEICNFSAEFLQNIIRLYN 180
PfSTONES      IFHQCLGLISQGNYIKSCRLLCPHMSILSLFSLYGTITFCSKEICNFSIELLQNIFRIFD 180
PvSTONES      ILQECLGLLSQGNYVKSCRMLSSHMGVLSIFSLLGTVTFCAKEICNFSVELLQNVLRIVN 180
PkSTONES      ILHQCLGLISQGNYVKSCRMLSSHMGVLSIFSLFGTVTFCAKEICNFSVELLQNVLRIVH 180
Consensus     *:::****:*****:****:*. **.:*::*** **:***:******* *:***::*: .

PbSTONES      IS-NEAKYGMTKIFVDLTTPG-AECASAPNTIYMESINYCVAFNDNVIVNFYENLWFIGN 238
PySTONES      IS-NEAKYGMTKVFVDLTNPG-KGCASDPNNIYMESINYCISYNDKVIVNFYENLWFIGN 238
PcSTONES      RS-EESKYGMTKIFVDFTTPG-SKCASAPNTIYMESINHCIEFNDHVIVNFYENLWFIGN 238
PfSTONES      RK-EEQKYGMTKIFIDLLNRD-IDCTYIPNSLYVESLGQCISYNDEVFVNFYENIWFISN 238
PvSTONES      MRNGEKKYAMTKVFIDLTKRHSEDCALMPDTIYLEAINQCVSFNRNVMVNFYENLWFIGN 240
PkSTONES      MENGEKKYGMTKVFIDLTKREMDDCSLMPDTIYLNSINQCISFNKNVMVNFYENLWFIGN 240
Consensus         * **.***:*:*: .     *:  *:.:*::::. *: :* .*:******:***.*

PbSTONES      KGNYIYLILISTLFILFCYFILLRENIRVFKFFIFVIIINWITTKSVVTMNTFFFYIP-Q 297
PySTONES      KGNYIYLILISTLFILFCYFILLRENIRVFKFFIFVIIINWITTKSVVTMNTFFFYIP-Q 297
PcSTONES      KDNYIYLILISTLFILFCYFILLRENIRVFKFFIFVIIINWITTKSVVTMNTFFFYIP-Q 297
PfSTONES      NKNYIYLIIISTLFLIFCYFLLLKENFRVFKFFIFVIIINWITTKSVVIMNTFFFYIPPQ 298
PvSTONES      KENCLYLILIATLFILFCYLILLRENFRVFKFFIFVIIVNWISSKSVVIMNTFFFYIP-Q 299
PkSTONES      KENYIYIILIATIFILFCYFILLRENFRVFKFFIFVIIVNWISSKSVVIMNTFFFYIP-Q 299
Consensus     : * :*:*:*:*:*::***::**:**:***********:***::**** ********* *

PbSTONES      SKKIYEEIISIFKEIGFLKLTIFLYGTLAR   IFILSTFASYFHTNTNIMTISLYTFVF 357
PySTONES      SKKIYEEIVSIFKEIGFLKLTIFLYGTLAR   IFILSTFASYFHTNTNIMTINLYTFGF 357
PcSTONES      PKKIYEEIVSIFEEIGFLKLTIFLYGTLAR   IFILSTFASYFHTNTNIMTISLYTFGF 357
PfSTONES      NQQIYEEILLIFKQVGYVKLTIFLYGTLAR   IFILSTFASYFNINTNIFKISIYTFAF 358
PvSTONES      EKKLHQEVVQIFQEIGIVKLTIFLYGTIAR   IFILSTFASYFHTNTNVLRITIYTFAF 359
PkSTONES      QIKLYQEVLEIFEEIGIIKLTIFLYGTIAR   IFILSTFASYFHTNTNILKITMYTFAF 359
Consensus       ::::*:: **:::* :*********:****************: ***:: *.:*** *

PbSTONES      YMLNMYFYLKNEAHEYYLYNKGLGS--GASFNFMKCNTYINMLQELKDNKGNI------- 408
PySTONES      YMLNMYFYLKNEAHEYFLYNKGLGS--GGKFNFMKCNTYINMLQELKDNNGNI------- 408
PcSTONES      YMMNMYFYLKNEAHEYYLYSKGLGS--GTSYNFMKCNTYINMLRELKNNNVNI------- 408
PfSTONES      YILNMYFYLKNEAYEYFLYNKSLSHNNTDEFNFIKCNTYINMLHEINNPNHNNHF----- 413
PvSTONES      YILNMYFYLKNEAHEYYLYNSRDF-STVKEFNFMKCNTYINLLQGLAHGRENGGAGGAAA 418
PkSTONES      YILNMYFYLKNEAHEYYLYNSKNL-ITSKEFNFIKCNTYINLLQGLAGDQRTEGKTDIPS 418
Consensus     *::**********:**:**..        .:**:*******:*: :   . .

PbSTONES      -------YNSGSSGSMYSVWQKLCLFFSSYSFLFSTVFIVLMQLLGPFNILTELAENYKY 461
PySTONES      -------YNSGSGSSMYSVWQKLCLFFSSYSFLFSTVFIVLMQLLGPFNILTEIAENYKY 461
PcSTONES      -------YNTASPSSMYSVWQKLCLFFSTYSFLFSTVFIVLMQLLGPFNILTEIAENYKC 461
PfSTONES      -------VYLYKYKQKYAVWQKLCLFFSTYSFLFSTIFIVLMQLLGPFNIINELIFTNNK 466
PvSTONES      SYREDDIPSAHTQDDRYAVWQKLCLFFSSYSFLFSTIFIVLMQLLGPFNILMERGRRDGH 478
PkSTONES      SYGV----THFTEDDKYAVWQKLCLFFSSYSFLFSTIFIVLMQLLGPFNILMEWRKSNSH 474
Consensus                .  . *:**********:*******:*************: *

(CONTINUES IN NEXT PAGE)

PbSTONES      -----NLLNTGKKIYTYKSLVKIYEKSDFKNISNSNRSYASSYYSKKKSVKSASDYEMAS 516
PySTONES      -----NLLNTGKKIYTYKSLVKIYEKSDFKNISNSNRSYTSSYYSKKKSVKSTSDYEIAS 516
PcSTONES      -----DLLDTGKKIYTYKSLVKIYEKSDFKNKSNSNRSYTSSYYSKKKSAKSASDYEMPS 516
PfSTONES      -----N-QQKDTKIYSYKSLVKKYEKYDIHHSLSTIIKS-----KRKRKSKNDGKYESKS 515
PvSTONES      VESEHSSGEGERRIYTYKSLVRRYERDRISRSASNNR   VSGRGGRAKRGRKNGYK--- 535
PkSTONES      ADEEDSNEERERKIYTYKSLVKRYERDRINRSVSNNRSSFLSKRGRSSTCSDKSTGR--- 531
Consensus          .  :   :**:*****: **:  : .  ..  .          .    .  .

PbSTONES      QKKK-KKKIFKNLNYFTTKPVKTIT---KNTESNITGKEKK------------EILIF-N 559
PySTONES      QKKK-KKNIFKNLNYFTSKPAKTMTT--KNTESNITGNEKKEKKEKKEKKEKKENFIFNT 573
PcSTONES      QKKKKKKKIFSNFNYFATKQAKTMTT--KNTESNITGKDKK------------EILDFNT 562
PfSTONES      KS-------------------------------KSNGKNKGED-------TNKFIHNYNT 537
PvSTONES      ---KGHKNIHQNGHKKGHQNGHKSSHQHGHKSSHQHGHKSSHQHGHK---NSHQHGHKNS 589
PkSTONES      ---RNPNGI---------------TH----------------------------IDRKNS 545
Consensus                                                                .

PbSTONES      TEEGGY-QTTDCAFSTN---K---------GVNLKKKK-EK---KESKMDFYNA------ 596
PySTONES      EEEGGY-QTADCAFSTN---K---------NVNLKNKK-EK---KGSKTDFYNA------ 610
PcSTONES      DEEGGY-QTTDCAFSNN---K---------GVSFKNKKPEK---KESKTNFYNV------ 600
PfSTONES      REDNYY-STYDKSDE--------------------------------HHDLAQI------ 558
PvSTONES      HQHGHKNSHQNCHPHCHQTDYGEKKKKFPRFLFFKGGEEAHRERHDQERSLYLSRKSSLL 649
PkSTONES      HK-----------------KKKRKKNFFRLNFFFKGRKVDRRGEENEERSLYLSRKSSFL 588
Consensus      :                                             . .:

PbSTONES      SKSNNFFDN------------------------------NNLKNSFSKKRQTNSSEYYCN 626
PySTONES      PQSGNLFGDNNNNNKNNNKNSNNKNNNNSNSNNNNSNSNTNLKSSFSKKRETNSSEYYCN 670
PcSTONES      PKSMNIFDN------------------------------EDLKNSFSKKRQTNSSEYYNN 630
PfSTONES      SNHM-----------------------------------NSIQS---SSYKKDSVSFYSS 580
PvSTONES      PEKGNYSQAHSSA-------------------------LSNLQNGG-EEAPDELLPYKCF 683
PkSTONES      PEKGDYFQEHS-A-------------------------FSNLQNGG-EI-QHDIPSYKSF 620
Consensus      :                                      .::.   .    :   :

PbSTONES      GKE--NKNIEESI-KDHN-KKNENNMLYSKNESSNDENSCMISTSTSTSNRESSYYEWTD 682
PySTONES      GKE--NQNIEGSI-KDDNNKKKEKNILYLKNE   DINSYIVSTSTSTSNRESSYCDWTD 727
PcSTONES      GKE--KKNIEGPI-KNDNNKK-KDNILYSKNESSNDENSGMIST   TSNRESSYYEWTD 686
PfSTONES      SH---STNME-II-KYDHGKKQ----TYDEK---NHSNSF   TNTDIDNNYNN------ 622
PvSTONES      DGEPGDFNFGGSSFEDDIGVAD----LAGEDLQGKKSNRRRRQSSTSA-HRKG   MYAD 738
PkSTONES      DRDPYDFNFGGSSFEDHIGVGD----LARENLQGKMTNRRQRQSSTSATQRNGSSNMYAD 676
Consensus     .    . *:     : .            :.   .  *    .:.:.  :. ..

PbSTONES      SDMSFGFMKNEKKDNNDWGMNKN-------------KGYQIGKEQN----NKTKNKNMNS 725
PySTONES      SDMSFGFMRNEKKDINDFGMDKN-------------KDDNIGKGEKYNGDDKKRSENMNR 774
PcSTONES      SDMSFGFMKNEKKDVNDWGMSKN-------------RNDQNEKDEQ----GTSKDINMNS 729
PfSTONES      ------IYKMEKKNIK----------------------------------------NMSD 636
PvSTONES      SE------------LPSWGARRDLLSA-SSGSSAGSSEAVEGGGARWSGEGASGKGGSGK 785
PkSTONES      SD------------PPSWGVRRRALFSDSSGPSASS--SDVGEGARWSGESRSRKENSGK 722
Consensus                                                             . .

PbSTONES      S-------------------------INEEYIVE--NKNKGMDIFRGNLFNRVINFISNI 758
PySTONES      N-------------------------MNEGYVVE--NKENGIYICRGNFIIRIKNFISKI 807
PcSTONES      N-------------------------INEKYILE--KKDKGMNRCRQNLFSRVINFISNI 762
PfSTONES      E-------------------------MMKKKTK---KDQKKKN-------------VVEN 655
PvSTONES      GISARGRPTKGSSPKGGSGRGDSGKSISAKTTSARGITGTVSPTRKNTLHRGIAALVEGM 845
PkSTONES      GSG-----------------------------------GTGVPKQKSDLHTGIARRVAKM 747
Consensus                                            .                :

PbSTONES      YNILLKKYGIFVLFFFIYIFTLMHIVDFKRNNNIVSEIILYNLYRSYLIFIIAAQIIYAS 818
PySTONES      YNILLKKYGIFVLFIFIYIFTIIHIVDFRRNDNIVSEIILYNLYRSYLIFIIAAQIIYAS 867
PcSTONES      YNILLKKYGIFVLFFFIYLFTLMHILDFKRNNNIVSEIILYNLYRSYLIFIIAAQIIYAS 822
PfSTONES      NLIRIKRYGNFFLFFIIYIFTILHIIDFKRNNNIISEIILYSMYRSFLIFIISCQIIYSS 715
PvSTONES      KNRVMNKYGTFLLFAVIYVFTILHVMDFRRNKDIVSEIILNSLYRGFLIFICAVQVIYAS 905
PkSTONES      KNLVINKYGTFLLFTIIYILTIFHVIDFKRNNDIVSEIILNSLYRGFLIFICAVQVIYAS 807
Consensus         :::** *.** .**::*::*::**:**.:*:***** .:**.:**** : *:**:*

(CONTINUES IN NEXT PAGE)

PbSTONES      WLFGMKMQMIQCGVISCLMNFITWIIILPLIFIIYSEKHTMIKLITIGIVLNIITIIISY 878
PySTONES      WLFGMKMQMIQCGVISCLLNFTTWIIILPLLFIIYSEKRNMIRLIIIGIVLNVITIIISY 927
PcSTONES      WIFGMKMQMIQCGVISCLMNFMTWIVILPLIFIVYSEEYTMTKLIIIGLFLNIITIIISY 882
PfSTONES      WIFGLRLQCSQCGIFSCLLHFITWIILLPGMFLIYMYKNYFIHIMIIGILLNIITIFISY 775
PvSTONES      WLFGMKLQMAQCGVISCLLNFFSWVVVLPLVFVIYFREGSWLQMLIIGLSLNVVAVATSF 965
PkSTONES      WVFGMKLQMSQCGVFSCLLNFFTWVVVLPLLSVLYLREGTWFQVFLIGLSLNAATVATSF 867
Consensus     *:**:::*  ***::***::* :*:::** : ::*  :    ::: **: **  ::  *:

PbSTONES      IEVIKKIKTQEKLCTSHKN---------KHDEMLQKFNKIVILKKCLYWLYIGNLEILRT 929
PySTONES      IEIIKKIKIQEKLCTSHKN---------KHDEMLQKFNKIIILKKCLYWLYIGNLEILRK 978
PcSTONES      IEVIKKIKIQEKLCTSHKN---------KHDEMIQKLNKIVIFKKCLYWLYIGNLEILRR 933
PfSTONES      IEVINKLKKNQQKITHRNNNIN-----TQYKKILYEYDKINILKKCIYWLYIGNIDILRK 830
PvSTONES      LEVTLKLRRGTA-------------------EKRQRYSYAALFKKCLYWLYIGNLEVLRR 1006
PkSTONES      VEVTVKLRRERASNISQKISMKEQGRVGHLAQNYQQYRYTTLFKKCLYWLYIGNLEVLRR 927
Consensus     :*:  *::                       :   .     ::***:*******:::**

PbSTONES      NLNIAMSGSEKKYIPFFWVVFFKYINSSMLIVIIIYNTKEYFFNMKDIFKATFSLKIELF 989
PySTONES      NLNIAMSGNEKKYIPFFWVIFFKYINSSMLIVIIIYNTREYFFNMKDIFKATFSLKIELF 1038
PcSTONES      NLNIAMSGSEKKYIPFFWIFFFKYINSSMLIVIIIYNTKEYFFNMKDVFRATFSLKVELF 993
PfSTONES      NMNIAISGNEKKYIPFFWVIFFKYINSSMLIVIIIVNTKEYFINMLYIFESSFSFNIELI 890
PvSTONES      NMNIAMSGQEEKYIPLFWSFFLKYINSSMLVVVVIYNTREYFFKMLPMFGGSFSFRIELG 1066
PkSTONES      NMNIAMSGNDEKYIPPFWTFFLKYINSSMLIVAVIYNTREYFFDMLPMFGCSFSLRIELI 987
Consensus     *:***:**.::**** ** .*:********:* :* **:***:.*  :*  :**:.:**

PbSTONES      SIFMIFFGSFWVLLRNIFKDHYMPQNIVIPSVPLGVHEKEKKKYIFPN 1037
PySTONES      SIFMIFFGSFCALVRNVFKDHYMPQNIVIPSVPLGVHEKEKKKYIFPN 1086
PcSTONES      SICMIFFGSFCVLVRNLLKDHYMPQNIVIPSIPLGVHEKERKRYIFPN 1041
PfSTONES      LILIIFIVSFFLLFKNIFTDNYNTQNVIIPPIPMVCQDK--KRYIFYE 936
PvSTONES      SIVLIFLGSLAVLAGNLRRDHYRPQTIGIPPVPMFCQDK--RRYIFAS 1112
PkSTONES      AILLIFLGSLAVLANNLRKDHYRPQGIIIPSVPMFCQDK--SRYIFSS 1033
Consensus      * :**: *:  *  *:  *:*  * : ** :*:  ::*   :*** .
```

##### Figure S3. Multiple sequence alignment of *Plasmodium* STONES orthologs.

PBANKA_0413500 (*STONES*) encodes a 1,037 amino acid (aa) protein (122 kDa) predicted to contain 14 transmembrane domains (aa 66-86, 106-129, 141-159, 242-261, 270-295, 319-340, 347-366, 420-442, 766-784, 804-823, 830-853, 859-881, 945-966 and 986-1003). Domain analysis revealed coupled, N-terminal TOF-LisH motifs in STONES, which are thought to promote oligomerization. *Stones* orthologs are present in all sequenced *Plasmodium* species, all predicted to encode 14 transmembrane domain proteins. Protein sequence alignment shows 88%, 92%, 64%, 59% and 61% similarity to *P. yoelii* (PY17X_0416300), *P. chabaudi* (PCHAS_0414400), *P. falciparum* (PF3D7_0315700). *P. vivax* (PVX_095450) and *P. knowlesi* (PKNH_0826400) STONES, respectively. Fully conserved amino acid residues are shaded in black and similar residues in grey. In the consensus sequence, identical residues among the orthologs are represented with an asterisk and the dots and colons mark amino acid residues with weakly and strongly similar amino acid characteristics. Protein sequences were retrieved from VeupathDB.

```
PbCRYSP      MKKGMQVYVLIYLIIFLEGYFSLSLFRTSVFHKEFTKTVERHLRDDYGDKDVAVFREIIR 60
PyCRYSP      MKKGMQVYVLIYLIIFLEGYFSLSLFSTSVFHKEFTKTVERHLREDYGDKDVAVFREIIR 60
PcCRYSP      MKKGMQVYVLIYLIIFLEGYFSLSLFSTSVFHKNFTKTVERHLREDYGDKDVAVFREIIK 60
PfCRYSP      MEKNVNVYLFLYLIIFLEAYFCVSLFSTNIFPREYTKVVEKHLREDYGDRDVEVFREIIR 60
PvCRYSP      MKRRTQGLLLVYLLTFLEAAFCLSLFSKSAFPREFTKSVERHLREDYGDRDVQVFREIMR 60
PkCRYSP      MKRRIPGLLLLFLFTFLEASFCLSLFSKSAFPREFTKAVERHLREDYGDRDVEVFREIMR 60
Consensus    *::     ::::*: ***. *.:*** .. * :::** **:***:****:** *****::

PbCRYSP      NYKNDDVFFNPSDEEKLKISIRKYAGDRFIQEYDHLMNEDSNDPQKVLAKSMISLIKQQF 120
PyCRYSP      NYKNNDVFLNPSDEENLKISMRKYAGDRFIQEYDNLMAENSNDSQKVLAKSMINLIKQQF 120
PcCRYSP      NYKNDDVFLSPSDEEKLKISMRKYAGDRFIQEYDNLMNENSNDSQKVLAKSMINLIKQQF 120
PfCRYSP      NYKDTDVFLSPSEEAKLKVNIQKYAGDHFIKEYENLMNEDTTDSNKKLAKTMINLIKQQF 120
PvCRYSP      NYKNEDTHLTPAEEEDLKAAMKKYAGSRFIQEYEALMDDNSKDSQKVLAKSMVNLIKQQF 120
PkCRYSP      NYKNEETHLTASDEDDLKVAMKKYAGNRFIQEYERLMDDNSKDSKKVLAKSMINLIKQQF 120
Consensus    ***: :..:. ::* .**  ::****.:**:**: ** :::.* :* ***:*:.******

PbCRYSP      IKLKEIETQYVTPNFEAYKEITKLKPLAQELDADTPCNTEAECKKLENMMNICTYVRSGA 180
PyCRYSP      IKLKEIETQYVTPNFEAYKEITKLKPLAQELDADTPCNTEAECKKLENMMNICTYVRSGA 180
PcCRYSP      IKLKEIETQYVTPNFEAYKEITKLKPLAQELDADTPCNTEAECKKLENMMNICTYVRSGA 180
PfCRYSP      IKLKVIEQEYITPNYEQYKQVAKLKPDISDLTADTPCNTEAECKKLENMMNICTYIRGGA 180
PvCRYSP      VKLKEIEAQYVTPNFDQYKQVAEMKPQLLDLNADTPCNTEAECKKLENMMNICTYVRGGA 180
PkCRYSP      VKLKEIEAQYVTPNFDQYKQVTELKPQVMDLNTDTPCKTEAECKKLENMMNICTYIRGGA 180
Consensus    :*** ** :*:***:: **:::::**   :* :****:*****************:*.**

PbCRYSP      DFAYDIFLVTTHVVTVMMAVLCACIFIGPVHVCALKNFPYTCKLPYPVFSTLFMATSSVW 240
PyCRYSP      DFAYDIFLVTTHVVTVMMAVLCACIFIGPVHVCALKNFPYTCKLPYPVFSTLFMATSSVW 240
PcCRYSP      DFAYDIFLVTTHVVTVMMAVLCACIFIGPVHVCALKNFPYTCKLPYPVFSTLFMATSSVW 240
PfCRYSP      DFAYDIFLVTTHVVTTMMAVMCACIFIGPVHICALKNFPYTCKLPYPIFSTLFMATSAVW 240
PvCRYSP      DFAYDIFLVTTHVVTSMMAVLCACIFIGPVHVCALKNFPYTCKLPYPVFSTLFMATSAVW 240
PkCRYSP      DFAYDIFLVTTHVVTSMMAVLCACIFIGPVHVCALKNFPYTCKLPYPVFSTLFMATSAVW 240
Consensus    *************** ****:**********:***************:*********:**

PbCRYSP      EVVKASTALCRVYGDLSIMSKMA 263
PyCRYSP      EVVKASTALCRVYGDLSIMSKMA 263
PcCRYSP      EVVKASTALCRVYGDLSIMSKMA 263
PfCRYSP      EVVKAATSLCRVYGDLSIMSKMA 263
PvCRYSP      EVVKAATALCRVYGDLSVMSKMA 263
PkCRYSP      EVVKAATALCRVYGDLSVMSKMS 263
Consensus    *****:*:*********:****:
```

##### Figure S4. Multiple sequence alignment of *Plasmodium* CRYSP orthologs.

*PBANKA_1338100 (CRYSP*) encodes 263 amino acid (aa) protein (30.4 kDa) predicted to contain three transmembrane domains (aa 6-26, 186-212, and 224-243) but no other domains. The protein is highly conserved amongst *Plasmodium* orthologs with 97%, 87%, 87%, and 86% sequence similarity to *P. yoelii* (PY17X_1342800), *P. chabaudi* (PCHAS_1342700), *P. falciparum* (PF3D7_1322900). *P. vivax* (PVX_116725) and *P. knowlesi* (PKNH_1205900) CRYSPs, respectively. PvCRYSP and PkCRYSP are predicted to encode only two transmembrane domains. Fully conserved amino acid residues are shaded in black and similar residues in grey. In the consensus sequence, identical residues among the orthologs are represented with an asterisk and the dots and colons mark amino acid residues with weakly and strongly similar amino acid characteristics. Protein sequences were retrieved from VeupathDB.

```
PbCRONE               1   MKEIIIFLFFFLYIACY-SIVLAKIPIENPPFIFSKNRKDDIN-MFLGNETISDD---LL 55
PyCRONE               1   MKEIITFLFFFLYIACY-TIVLAKIPIENPPFIFSKNRKDDIN-MFLGNETINDD---LL 55
PcCRONE               1   MKEIIIFLFFFLYIACY-SIVLSKIPIENPPFIFSKNSKDDIN-MFLGNETISDD---LL 55
PfCRONE               1   ---MNCVFFLCLYIFLG-SVVHLKVPIENAPFVFNK-KENDIK-YFLDKETPDDF---LS 51
PvCRONE               1   MSRVNFFALPLICFLLHGLAALAKIAIENPPFVFGNSGEDKLKRAFLNVDTPWDNDDVLS 60
PkCRONE               1   MSRVHFFVLPLLCFLLHGLAALAKIPIENPPFVFTNSNQDKLKRNFLNVDTPWDNDDVLA 60
Consensus                    :  . :  : :      .  *:.***.**:* :  ::.::  **. :*  *    *

PbCRONE               56  KPSYLMDVTLENFPNPFIHPSLCNRNGLKYSYICDPNKILSTYTADKIEEILSYQRRNSS 115
PyCRONE               56  KPSYSMDVTLENFPNPFIHPSLCNRNGLKYSYICDPNKILSTYTADKIEEILSYQRRNSS 115
PcCRONE               56  RPSYSMDITLENFPNPFIHPSLCNRNGLKYSYLCDPNKILSTYTADKIEEILSYQRRNSS 115
PfCRONE               52  NNSYSLDITLENFPNPFFHPSLCNRYGFKFSYICDPNKILSRNIADQIEEILNYQRRNSK 111
PvCRONE               61  KPPYSLYVTLENFPNPFIHASLCNRNGLNYSYVCDPNKILSRSTADKLEEILSYQRRNSS 120
PkCRONE               61  KPPYSLYVTLENFPNPFIHPSLCNRNGLSYSYLCDPNKILSRSTADKIEEILSYQRRNSS 120
Consensus                . .* : :*********:*.***** *:.:**:********   **::****.******.

PbCRONE               116 HYCITKGKVPYVLGVALVKKLPYGISADTFGSHILEYWRLGNTSCNDGILLLFVKDDINF 175
PyCRONE               116 HYCIAKGKVPYVLGVALVKKLPYGISADIFGSQILEYWRLGNTNCNDGILLLFVKDDINF 175
PcCRONE               116 HYCIDKGKVPYVLGVALVKKLPYGISADTFGSQILEHWRLGNKNCNDGILLLFVKDDINF 175
PfCRONE               112 HFCVDK-EVPYVLGVALINKLPYGISADTFASQIFEYWKLSNKDCNDGVLLFFVKEDTHF 170
PvCRONE               121 HHCADRGEVPYVLGVALIERLPYGVSAETFSSQILEHWKLGNRNCNDGILLLFVKDDATF 180
PkCRONE               121 HYCADRGEVPYVLGVALIERLPYGVSAETFSSQILEHWKLGNRNCNDGILLLFVKENATF 180
Consensus                *.*  : :*********:::****:**: *.*:*:*:*:*.* .****:**:***::  *

PbCRONE               176 VLKWKKGAQSIINFRTASSMNKTFKQYIRRYSLEYSILRAVKLTSQYLTEEIIPSTQTAQ 235
PyCRONE               176 VLKWKKGSQSIINFRTASSMNKTFKQYIRRYSLEYSILRAVKLTSQYLTEEIIPSTQTAQ 235
PcCRONE               176 VLKWKKGSQSIINFRTASSMNKSFKQYIRRYSLEYSILKAVKLTSQYLTEEIIPSTQTAQ 235
PfCRONE               171 ILKWKKGAQSIINFRTATSMNKSFNQYLRKYSLEYSILSAVKLTSQYLTEEIIPPTQTAQ 230
PvCRONE               181 VLKWRKGAQSIINFRTATAMNKSFNMYIRRYSLEYSILRAVTLTSQYLTEEIIPPTQTAQ 240
PkCRONE               181 VLKWRKGAQSIINFRTATAMNKSFNMYIRRYSLEYSILRAVTLTSQYLTEEIIPPTQTAQ 240
Consensus                :***:**:*********::***:*: *:*:******** **.************.*****

PbCRONE              236 MVVALTIAVIVGLSYLACILIVFSDAQKAN- 265
PyCRONE              236 MVVALTIAVIVGLSYLACILIVFSDAQKAN- 265
PcCRONE              236 MVVALTIAVIVGLSYLACILIVFSDAQKAN- 265
PfCRONE              231 KVVAFTIVIVVGLAYVAFILIVFADAQRDNK 261
PvCRONE              241 IVVALTIGIVVGLGYLACILIVFSDAQKGNI 271
PkCRONE              241 MVVALTIGIVVGLGYLACILIVFSDAQKGNR 271
Consensus                 ***:** ::***.*:* *****:***: *
```

##### Figure S5. Multiple sequence alignment of *Plasmodium* CRONE orthologs.

*PBANKA_0720900 (CRONE)* encodes a 265 amino acid (aa) protein (30 kDa) with an N-terminal signal peptide (aa 1-23), a C-terminal single-pass type I transmembrane domain (aa 236-258) and a TPM domain (PFAM04536; aa 130-172). This protein architecture is predicted for all *Plasmodium* orthologs. The TPM domain, named after its founding proteins TLP18.3, Psb32 and MOLO-1, the first two of *Arabidopsis thaliana* and the third of *C. elegans*, respectively (Sirpiö et al., 2007, Wegener et al., 2011, Wu et al., 2011, Boulin et al., 2012), despite a strong structural conservation exhibits a very low sequence conservation (Eletsky et al., 2012). Nevertheless, CRONE shows high sequence similarity with its *Plasmodium* orthologs ranging from 97% with *P. yoelii* (PY17X_0720900) and 97% with *P. chabaudi* (PCHAS_0729900) to 77%, 78% and 80% with *P. falciparum* (PF3D7_0418800), *P. vivax* (PVX_090030) and *P. knowlesi* (PKNH_0511500) CRONE, respectively. Fully conserved amino acid residues are shaded in black and similar residues in grey. In the consensus sequence, identical residues among the orthologs are represented with an asterisk and the dots and colons mark amino acid residues with weakly and strongly similar amino acid characteristics. Protein sequences were retrieved from VEuPathDB.

```
PbROVER    MHILQEDTNHFNSEHSGFGNQFINFSESYHKIAGEENYKNSKLNEIIAHNIYDNIKNNIN 60
PyROVER    MQILQEDTNHFNSEHSGFGNQFINFSESYHKIAGEENYKNSELNEIIAHNIYDNIKNNIN 60
PcROVER    MHILQEDTNHFNSEHSGFGNQFINFSEPYHKIAGEENYKNSELNEIIAHSIYDNIKSNIN 60
PfROVER    ----------------------MYFSESYQKIAGRKNYKKNALDSVSSQGIFDNIKNNIY 38
PvROVER    -----------------------MLSVPYHKIAKQERLEDPELKRVLAQNIYDNVKNNIA 37
PkROVER    -----------MPERSGCDTNFIHFSESYHKITKQEQFEDPELKRVLAQNIYDNLKNSIA 49
Consensus                          :*  *:**: .:. :.  *. : ::.*:**:*..*

PbROVER       YIKETQYNSCQDISTINSD------N----NHSDKN--------------KEGKHNYECL 96
PyROVER       YIKETQYNSCQDISTTNSN------N----NHFVKN--------------KEGKHNCEYL 96
PcROVER       YIKETQYNSCQDISPNNSD------N----SHFVKN--------------KEGKRNCECL 96
PfROVER       NIDKTDYKILNHYNYKKDQTSTNLSKNKQKDHFFIRSNNINNNNINNNNNNKGKDMCV-- 96
PvROVER       YLEETQYNPCECNHCRSDIADAPLED----TELRINVST------AKLGYKRESEKACVE 87
PkROVER       YLEETQYNPCECNHCRSDIADVPLED----TELLKNVPT------ANLIFKEESAKCSVE 99
Consensus      :.:*:*:  :     ..       .     .   .              :. .

PbROVER       NNNISLR--NNIK----FTNSKADSHNETHNILKNDINNLEIHQYIPYISSTEFLE---Q 147
PyROVER       NNSISLR--NNIK----FTNSKEDSHNESHNILKNDINNLGIHQCIPYISSTEFLE---Q 147
PcROVER       DNNISVR--NNIK----FTNSKADSHNETHNILKNDINNLEIHQCIPYITFTEFLE---Q 147
PfROVER       NPPYEKNILYYINNKNLGRYKNSCDHPTVHNILKNNKSNIEIHQYIPSLTSTIFDLLYVN 156
PvROVER       DREITQNNVQLIS----EVEGETSLHPKTYNILKNHKDNLEIHQSIPPLKCESLAEFTPR 143
PkROVER       DTKITPNNLSNIN----PVEGETSVHSKTYNILKNQKDNLEIHQIIPPLMYDSFSEFTPR 155
Consensus     :     .    *.        :   *   :*****. .*: *** ** :    :     .

PbROVER       NKKSKEINNLYQ----TYETMNDPRYNDYPPMNGFG--YNHNNMSNQNGFYTNTYNKELA 201
PyROVER       SKKSKEINNLYP----TYKTMNDPRYNDYPPMNGFG--YNHHNTSNENGFYTNTYNKELA 201
PcROVER       N---NEINNLYQ----EHKTMNDPRYNDYPPMNGFG--YNHNNTSNGNGFYTNTYNKELP 198
PfROVER       SNKIE--KNIYPQQNNNKNMSHSQKFNEYIPMNGFRSIYHNNNMNNGNGYYTNTFNKHNS 214
PvROVER       RSPHQKSQRLPK----EEDKMNNHRYGEYSPMNGFG--YNHHNSMNANGFCTNTDNREMA 197
PkROVER       SDSHHKSPSLPK----EEDKMNNNRYGEYSPMDGFG--YNHRNSMDANGFSTNTYNRELP 209
Consensus         .    :        .  :. ::.:* **:**   *::.*  : **: *** *:.

PbROVER       NLTMQQHNMHGRNNLMFEDGNIPYGLYDQSNYTGTNSVMCGNG-ISNEPKMMYRRNEMNN 260
PyROVER       NLTMQQHNMHGRNNLMFEDGNIPYGLYDQSNYTGTNSVMCGNG-VANEPKMMYRRNEMNS 260
PcROVER       NLAMQQHNMHGRNNLMFEDGNIPYGLYDQSNYTGTNSVMCGNG-VSNDPKMMYRRNEMNS 257
PfROVER       NLYTHQNNMYGRNNLMFDEDSIPYGMYDQSNITGTNTMTCGKN-IPCDERMMYRYNTMEG 273
PvROVER       NLTMQQHNMHGRNNLMFEDGNIPYGLYGQANYTGTNSLMCGGTPPAYDPKMMYRRNHETN 257
PkROVER       NLTMQQHNMHGRNNLMFEDGNIPYGLYGQSNYTGTNSLTYGGVNPSHDPKAMYRRNHETN 269
Consensus     **  :*:**:*******::..****:*.*:* ****::  *      : : *** *   .

PbROVER       P-PYYHNYP-MNNGMGYNEHIVPAMQYNPNMRKHMMRNGYSHECNYMNSRPNY-----PT 313
PyROVER       P-PYYHNYP-MNNGMGYNEPIVPPMQYNPNMRKNMMRNGYSHECNYMNSRPNY-----QI 313
PcROVER       P-PYYPNYP-MNNGMGYNEPIMPAMQYNPNMRKNMMRNAYSHECNYMNSRPNY-----SG 310
PfROVER       VGPCHNIHPMINTRMNYSTPLSLP-QYNPNIRNYLTHEGYPPQYYDYNSNLYSRNKRVLK 332
PvROVER       E-PFYNHYP-VASGMGYNVPVLPPQPYYPSMKRNLTKEGYTHGYHYMNNASSH----GAL 311
PkROVER       E-PFYNNYP-MGSGMGYNTPFLPPQAYYPNMKRNLTKEGYTHGCHYMNNPSNN----GAL 323
Consensus       * :  :* : . *.*.  .     * *.::. : ::.*       *.

PbROVER       PKNN-MDITSEKNYKKDNCRKNISKIKKGDIDINIETSSSSKKGVIIDLNIGVGN 367
PyROVER       PKNN-MDMTSERNYKKDNCRKNISKIKKGDIDINIETSSSSKKGVIIDLNIGVGN 367
PcROVER       PKNN-MDNTSEKSYKKDNCRKNISKIKKGDIDINIETSSSSKKGVIIDLNIGVGN 364
PfROVER       QSYDADNIDNEKNHKKELSRKNISKIKKGDIDINIETSSSDKRGVVIDLNIGV-- 385
PvROVER       PKYP-SDSVTDKYPKKESCRKNVSKVKKGDIDINIETSSSGRKGVIIDLNIGVGG 365
PkROVER       SKYP-SESVADKSPKKESCRKNLSKVKKGDIDINIETSSSSRKGVIIDLNIGMGN 377
Consensus      .    :   ::  **: .***:**:**************.::**:******:
```

##### Figure S6. Multiple sequence alignment of *Plasmodium* ROVER orthologs.

*PBANKA_1353800* (*ROVER*) encodes a 367 amino acid protein (43 kDa) with no signal peptide, transmembrane domain, or any other functional domain. Multiple sequence showed that PbROVER show very high sequence similarity with its *P. yoelii* (PY17X_1359200) and *P. chabaudi* (PCHAS_1358400) orthologs of 93% and 91%, respectively. Lower sequence similarity of 48%, 54% and 58% is seen with its *P. falciparum* (PF3D7_1340400), *P. vivax* (PVX_082937) and *P. knowlesi* (PKNH_1261000) orthologs, respectively. Identical amino acid residues are shaded in black and similar residues in grey. In the consensus sequence, identical residues among the orthologs are shown with an asterisk, and dots and colons mark residues with weakly and strongly similar characteristics. Protein sequences were retrieved from VEuPathDB.

```
PbSPM1         1   ---MEIIGAKPSVNFNFFDEETQNNDN-LNYMESTNNYDKREDQDYFYSNRRTFLDKNSE 56
PySPM1         1   ---MEIIGAKPSVNFNFFDEETQNNDN-LNYLESKNNYDKREDQDYFYSNRRMFLDKNSE 56
PfSPM1         1   ---MEVITEKPKVQFNFYDEQNNSYNNSTDYPEFSKDSN-IDYQPYVYTNRKFPLDKNSE 56
PvSPM1         1   ---MEIIAEKPKVKFNFASEEYKNCDS-SDYSECAEDYGRPNGKDYFYANRILSLDRNSE 56
PkSPM1         1   ---MEIIAEKPKVKFNFVPEDYKNCDN-SDYSECAEDYGRPNEKDYFYANRILSLDRNSE 56
Consensus             **:*  **.*:***  *: :. :.  :* *  :: .  : : * *:**   **:***

PbSPM1         57  NRRKESPSKKLGLCVDEICTCGFHRCPKIIKPLPFEGESNYRSEFGPKPLPELPPRVETK 116
PySPM1         57  NRRKESPSKKPGLCVDEICTCGFHRCPKIIKPVPFEGESNYRSEFGPKPLPEIQPRIENK 116
PfSPM1         57  LRRKESPSRKPGLCVDEICTCGFHRCPRIIKSIPFEGESNYRSEFGPKALPELPPQIYMK 116
PvSPM1         57  QRRKESPSKRPGLCVDEICTCGFHRCPKIVKSLPFDGESNYRSEFGPKPLPELPPRQEAK 116
PkSPM1         57  QRRKESPSKRPGLCVDEICTCGFHKCPKIVKPLPFEGESNYRSEFGPKPLPELPPRQDAK 116
Consensus           *******:: *************:**:*:*.:**:************.***: *:   *

PbSPM1         117 LVKSLPFEGESNYRSEFGPKPLPEIQPRVENKPHKTLPFEGESNYRSEFGPKPLPELPPR 176
PySPM1         117 PHRTLPFEGESNYRSEFGPKPLPEFPPRVETKLVRSLPFEGESYYRSEFGPKPLPELPPR 176
PfSPM1         117 PPKPLPFEGQTNYRSEFGPKPLPELPPQIYMKPPKSLPFEGETNYRSEFGPKPLPELPPR 176
PvSPM1         117 LTRSLPFEGESNYRSEFGPKPLPELPPRVEQKPPKSLPFDGESNYRSEFGPKPLPELPPR 176
PkSPM1         117 LMRSLPFEGESNYRSEFGPKPLPELPPRVEQKVPKSLPFEGESNYRSEFGPKPLPPLPPR 176
Consensus            :.*****::*************: *::  *  ::***:**: *********** ****

PbSPM1         177 VETKLVKSLPFEGE--------------------------------SNYRSEFGPKPLPE 204
PySPM1         177 IVTKLVKSLPFEGESYYRSEFGPKPLPEIQPRVENKPHKTLPFEGESNYRSEFGPKPLPE 236
PcSPM1         180 VVTKLVKSLPFEGE--------------------------------SYYRSEFGPKPLPE 207
PfSPM1         177 IETKLVKPLPFEGESNYRSEFGPKPLPELPPQIYMKPPKSLPFEGETNYRSEFGPKPLPE 236
PvSPM1         177 VEQKPPKSLPFDGESNYRSEFGPKPLPELPPRVEQKPPKSLPFEGESNYRSEFGPKPLPE 236
PkSPM1         177 VVTKLVKSLPFEGESNYRSEFGPKPLPELPPRVEQKAPKSLPFEGESNYRSEFGPKPLPE 236
Consensus          :  *  *.***:**                                : ************

PbSPM1         205 IQPRVENKPHKTLPFEGES----------------------------------------- 223
PySPM1         237 IQPRVENKPHRTLPFEGES----------------------------------------- 255
PcSPM1         208 IQPRVEHKPHKTLPFEGES----------------------------------------- 226
PfSPM1         237 LPPQIYMKPPKSLPFEGETNYRSEFGPKPLPELPPRIETKLVKPLPFEGESNYRSEFGPK 296
PvSPM1         237 LPPRVEQKPPKSLPFEGESNYRSEFGPKALPELPPRVEQKPPKSLPFEGESNYRSEFGPK 296
PkSPM1         237 LPPRVEQKAPKSLPFEGES----------------------------------------- 255
Consensus          : *::  *. ::******:

PbSPM1         223 -------------------------------------------------------NYRSE 228
PySPM1         255 -------------------------------------------------------NYRSE 260
PcSPM1         226 -------------------------------------------------------NYRSE 231
PfSPM1         297 PLPELPPQIYMKPPKSLPFEGETNYRSEFGPKPLPELPPQIYMKPPKQLPFEGESNYRSD 356
PvSPM1         297 PLPALPPRVETKLVKSLPFEGESNYRSEFGPKPLPELPPRVEQKPPKSLPFEGESNYRSE 356
PkSPM1         255 -------------------------------------------------------NYRSE 260
Consensus                                                                 ****:
```

MAP6 domain

```
PbSPM1         229 FGPKPLPELPPRVETKLVKSLPFEGESYYRSEFGPKPLPEIQPRVENKPHKTLPFEGES- 287
PySPM1         261 FGPKPLPELPPRVETKLVKSLPFEGESHYRSEFGPKPLPEMQPRVENKPHKTLPFEGESN 320
PcSPM1         232 FGPKPLPELPPRVETKLVKSLPFEGESHYRSEFGPKPLPEIQPRIENKPPKSLPFEGESN 291
PfSPM1         357 YGPKPLPELPPRIEMKLPKSLPFEGESNYRSEFGPKPLPELPPKIYMQPPKPLPFEGESN 416
PvSPM1         357 FGPKPLPALPPRVVTKLVKSLPFEGESNYRSEFGPKPLPEIPPRVEQKPPKSLPFEGESN 416
PkSPM1         261 FGPKPLPPLPPRVVTKLVKSLPFEGESNYRSEFGPKPLPELPPRVEQKAPKSLPFEGESN 320
Consensus          :****** ****:  ** ********* ************: *::  :. *.*******

PbSPM1         287 ------------------------------------------------------------ 287
PySPM1         321 YRSEFGPKPLPEIQP--------------------------------RIENKPHRTLPFE 348
PcSPM1         292 YRSEFGPKPLPELPP--------------------------------RVENKPHKTLPFE 319
PfSPM1         417 YRSEFGPKPLPELPP--------------------------------RHETKLVKQLPFE 444
PvSPM1         417 YRSEFGPKPLPELPPRVEQKPPKSLPFEGESNYRSEFGPKQLPELPPRQEAKLTRSLPFE 476
PkSPM1         321 YRSEFGPKPLPELPP--------------------------------RQETKLTRSLPFE 348
Consensus

(CONTINUES IN NEXT PAGE)

PbSPM1         287 ---NYRTEYVRKVIPICPVNLLPKYPAPTYPAEHIFWDDVKKAWY 329
PySPM1         349 GESNYRSEYVRKVIPICPVNLLPKYPTPTYPSEHIFWDDVTKAWY 393
PcSPM1         320 GESNYRSEYVRKVIPICPVNLLPKYPTPTYPSEHVFWDDVKQTWY 364
PfSPM1         445 GESSYRTEYIRKVLPVCPVELLPKYPTPTYPSQHVFWDRETKKWY 489
PvSPM1         477 GESSYRSEYVRKAIPICPVNLLPKYPAPTYPSEHVFWDSACKRWY 521
PkSPM1         349 GESSYRSEYVRKAIPICPVNLLPKYPAPTYPSEHVFWDSTCKRWY 393
Consensus             .**:**:**.:*:***:******:****::*:***   : **
```

##### Figure S7. Multiple sequence alignment of *Plasmodium* SPM1 orthologs.

*PBANKA_0810700* (*SPM1*) encodes a 329 aa (38 kDa) putative subpellicular microtubule protein, SPM1, predicted to contain a microtubule associated protein 6 (MAP6) domain (IPR024963) in aa 69-286 and no signal peptide or transmembrane domain. Subpellicular microtubules are a basic scaffold that consists of a row of microtubules that run under the pellicle and determine the rigid cell shape of several protists. They originate from the apical polar ring and their interaction with the cytosolic face of the pellicle is mediated by the microtubule associated proteins (MAPs), including SPM1 and its paralog SPM2 originally identified in *Toxoplasma gondii* (Tran et al, 2012). SPM1 homologs are also present in other apicomplexan parasites (Tran et al, 2012). SPM1 protein comparisons with other *Plasmodium* SPM1 orthologs revealed a similarity of 78%, 84%, 55%, 56% and 72% with *P. yoelii* (PY17X_0814000), *P. chabaudi* (PCHAS_0811000), *P. falciparum* (PF3D7_0909500). *P. vivax* (PVX_098915) and *P. knowlesi* (PKNH_ PKNH_0707500) SPM1 orthologs, respectively*.* This analysis also showed that within the MAP6 domain, PbSPM1 is made up of seven 26 aa repeat motifs. These repeats are conserved within *Plasmodium* orthologs, ranging in number from 8 in PySPM1 to 13 in PvSPM1. Fully conserved amino acid residues are shaded in black and similar residues in grey. In the consensus sequence, identical residues among the orthologs are represented with an asterisk and the dots and colons mark amino acid residues with weakly and strongly similar amino acid characteristics. Protein sequences were retrieved from VEupathDB.

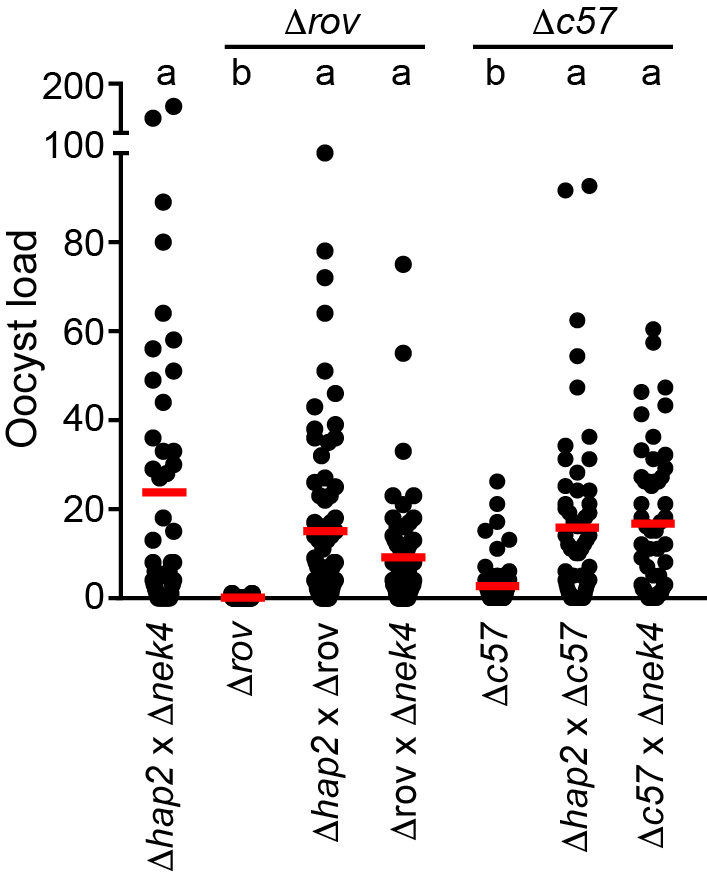

##### Figure S8. Genetic complementation assays of *rover* and *pimms57*.

Oocyst distribution in the midguts of *A. coluzzii* mosquitoes at day 9 pbf, following genetic crossing of *Δrov* or *Δc57* with either *Δnek4* or *Δhap2* lines, respectively. Infections of the *c507 wt* line and a genetic cross between *Δnek4* and *Δhap2* were used as controls. Two biological replicates were performed and statistical analysis was performed using a Mann–Whitney U-test. Letters (a, b and c) above each infection indicate the results from multiple comparisons: no difference was detected between infections annotated with the same letter, while a statistically significant difference (p < 0.0001) was detected between infections annotated with difference letters. The mean is shown with a red line.

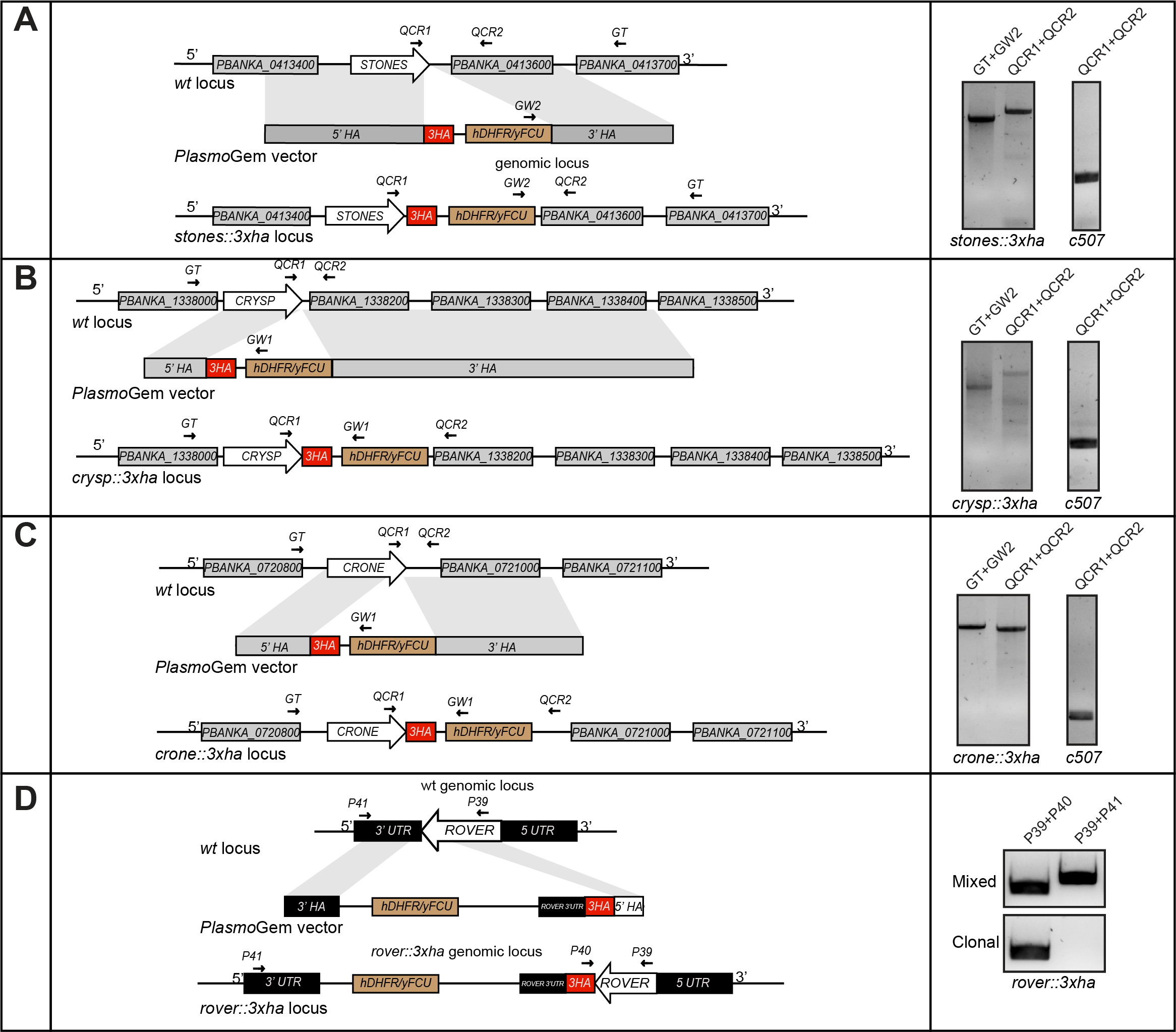

##### Figure S9. Generation of gene tagged parasites.

Schematic representation of the 3XHA tagging of (**A**) *STONES*, (**B**) *CRYSP* (**C**) *CRONE* and (**D**) *ROVER* in the *c507* line by using PlasmoGEM gene tagging vectors that carry the *hDHFR)/yFCU* selectable marker. In each panel, the *wt* genomic locus (top), the plasmid carrying the homologous arms (HA), the transfected DNA fragment carrying the tagging cassettes and the selectable markers (middle), and the final transgenic locus after double crossover homologous recombination (bottom) are shown. Constructs are not drawn to scale. Small arrows show the primers used for the confirmation of integration of gene targeting constructs and deletion of the gene of interest using diagnostic PCR reactions as shown in the panels on the right.

### Supplementary Tables

**Table S1. *P. berghei* gametocyte expressed genes in the *A. coluzzii* midgut 1 hour post infection.**

**(A)** The data are presented as an average of log2-transformed signal intensity ratios of normalized ANKA 2.34 vs. ANKA 2.33 infected midguts. The expression data for 3428 analyzed genes are shown. P-value was calculated using a one-way ANOVA following correction with the Benjamini-Hochberg hypergeometric. *P. berghei* PlasmoDB gene ID, gene names and product description are also given. 189 statistically significant up-regulated genes **(B)** and their domain analysis **(C)** are presented. Differentially regulated expression data in DOZI and CITH knockout mutants are presented for the 189 genes. *P. berghei* single cell transcriptomic raw count data of male and female gametocytes is also presented. PlasmoDB gene and functional annotation is also shown.

**Table S2. PlasmoGEM barcode sequence counts.**

Barcode counts shown are raw counts normalized to 1000 counts per sample. Ooc is oocysts and spz is sporozoites. Assays were performed in 3-4 replicates designated as r1-r4.

| **Table S3. Oocyst counts of mutant parasites and *c507* controls in *A. coluzzii*** | | | | | | | |
| --- | --- | --- | --- | --- | --- | --- | --- |
| Parasite | Experiment | No of  midguts | Prevalence  (%) | Arithmetic  mean | Median | Parasite  Range | P value |
| *c507* | Pooled | 61 | 82 | 22.1 | 13 | 0-176 |  |
| *Δsto* |  | 67 | 0 | 0 | 0 | 0 | <0.0001 |
| *c507* |  | 80 | 86 | 24.6 | 14 | 0-127 |  |
| *Δcry* |  | 76 | 93 | 40.1 | 21 | 0-238 | 0.0518 |
| *c507* |  | 99 | 89 | 70.9 | 47 | 0-400 |  |
| *Δcro* |  | 99 | 87 | 56.1 | 33 | 0-277 | 0.4245 |
| *c507* |  | 120 | 78 | 28.4 | 9 | 0-203 |  |
| *Δrov* |  | 104 | 5 | 0.05 | 0 | 0-1 | <0.0001 |
| *c507* |  | 106 | 85 | 39.3 | 32 | 0-176 |  |
| *Δspm1* |  | 105 | 88 | 27.5 | 15 | 0-136 | 0.0535 |
| *Δsto* | R1 | 32 | 0 | 0 | 0 | 0 | <0.0001 |
| *c507* |  | 26 | 81 | 28.3 | 16 | 0-120 |  |
| *Δcry* |  | 26 | 88 | 58.1 | 41 | 0-238 | 0.0579 |
| *c507* |  | 34 | 94 | 122.0 | 95 | 0-400 |  |
| *Δcro* |  | 34 | 94 | 91.9 | 75 | 0-277 | 0.1813 |
| *c507* |  | 40 | 85 | 29.7 | 6 | 0-203 |  |
| *Δrov* |  | 33 | 9 | 0.1 | 0 | 0-1 | <0.0001 |
| *c507* |  | 34 | 74 | 34.8 | 16 | 0-176 |  |
| *Δspm1* |  | 35 | 91 | 26.0 | 18 | 0-103 | 0.9166 |
| *Δsto* | R2 | 35 | 0 | 0 | 0 | 0 | <0.0001 |
| *c507* |  | 28 | 93 | 24.8 | 17 | 0-127 |  |
| *Δcry* |  | 25 | 96 | 26.9 | 16 | 0-208 | 0.5505 |
| *c507* |  | 35 | 80 | 29.3 | 9 | 0-218 |  |
| *Δcro* |  | 35 | 74 | 25.1 | 11 | 0-133 | 0.9930 |
| *c507* |  | 35 | 60 | 26.0 | 3 | 0-166 |  |
| *Δrov* |  | 35 | 0 | 0 | 0 | 0 | <0.0001 |
| *c507* |  | 32 | 94 | 48.7 | 44 | 0-149 |  |
| *Δspm1* |  | 30 | 90 | 27.9 | 18 | 0-118 | 0.0422 |
| *c507* | R3 | 26 | 85 | 20.6 | 9 | 0-110 |  |
| *Δcry* |  | 25 | 96 | 34.6 | 24 | 0-105 | 0.0818 |
| *c507* |  | 45 | 84 | 29.2 | 15 | 0-137 |  |
| *Δcro* |  | 30 | 93 | 51.7 | 33 | 0-165 | 0.8862 |
| *c507* |  | 40 | 88 | 35.7 | 29 | 0-158 |  |
| *Δrov* |  | 36 | 6 | 0.1 | 0 | 0-1 | <0.0001 |
| *c507* |  | 30 | 93 | 61.5 | 44 | 0-276 |  |
| *Δspm1* |  | 40 | 83 | 28.7 | 8 | 0-136 | 0.1883 |
| *c507* |  | 40 | 88 | 35.7 | 29 | 0-158 |  |

| **Table S4. Oocyst counts of mutant parasites and *c507* controls in *A. coluzzii*** | | | | | | | | |
| --- | --- | --- | --- | --- | --- | --- | --- | --- |
| Genetic crosses | Experiment | | No of  midguts | Prevalence  (%) | Arithmetic  mean | Median | Parasite  Range | P value |
| *Δhap2 x Δnek4* | | pool | 47 | 79 | 23.7 | 11.8 | 0-154 |  |
| *Δrov* | |  | 55 | 11 | 0.1 | 0.05 | 0-1 | <0.0001 |
| *Δhap2 x Δrov* | |  | 76 | 84 | 15 | 7.5 | 0-38 | 0.59 |
| *Δnek4 x Δrov* | |  | 58 | 74 | 9.1 | 4.6 | 0-75 | 0.59 |
| *Δc57* | |  | 69 | 49 | 2.6 | 1.3 | 0-26 | <0.0001 |
| *Δhap2 x Δc57* | |  | 58 | 79 | 15.7 | 7.9 | 0-92 | 0.64 |
| *Δ nek4 x Δc57* | |  | 53 | 87 | 16.6 | 8.3 | 0-60 | 0.98 |

| **Table S5. Midgut and salivary glands sporozoites and mosquito-to-mouse transmission** | | | | | | | | | |
| --- | --- | --- | --- | --- | --- | --- | --- | --- | --- |
|  |  | Midgut sporozoites | | | Salivary gland sporozoites | | | |  |
| Parasite | Reps | Mean | SEM | P | Reps | Mean | SEM | P | Bite-back |
| *c507* | 2 | 3,567 | 259 |  | 2 | 1,843 | 83 |  | 6/6(3/3;3/3) |
| *Δsto* | 2 | 10 | 7 | 0.011 | 2 | 67 | 0 | 0.004 | 0/6(0/3;0/3) |
| *c507* | 3 | 6,036 | 180 |  | 3 | 3,484 | 480 |  | 6/6(2/2;2/2;2/2) |
| *Δcry* | 3 | 6,391 | 745 | 0.99 | 3 | 0 | 0 | 0.004 | 0/6(0/2;0/2;0/2) |
| *c507* | 3 | 10,694 | 902 |  | 3 | 7,587 | 925 |  | 9/9(3/3;3/3;3/3) |
| *Δcro* | 3 | 0 | 0 | 0.001 | 3 | 0 | 0 | 0.003 | 0/9(0/3;0/3;0/3) |
| *c507* | 3 | 3,556 | 970 |  | 3 | 4,344 | 1,086 |  | 9/9(3/3;3/3;3/3) |
| *Δrov* | 3 | 61 | 11 | 0.042 | 3 | 16 | 9 | 0.031 | 0/9(0/3;0/3;0/3) |
| *c507* | 3 | 2,268 | 488 |  | 3 | 4,668 | 458 |  | 9/9(3/3;3/3;3/3) |
| *Δspm1* | 3 | 1,121 | 206 | 0.152 | 3 | 2,007 | 28 | 0.009 | 9/9(3/3;3/3;3/3) |
| For each biological replicate, sporozoite numbers was determined from 25-30 homogenized mosquito midguts or salivary glands at 15 and 21 dpbf respectively. Infectivity of sporozoites was assessed by infected mosquito bite back experiments with at least 30 mosquitoes on C57/BL6 mice at 21 dpbf. Following this, parasitemia was monitored until 14 days post mosquito bite. *P* values were calculated using the unpaired Student’s t-test. SEM represents standard error of mean. | | | | | | | | | |

| **Table S6. Invasion assay in *CTL4* knockdown *A. coluzzii*** | | | | | | | |
| --- | --- | --- | --- | --- | --- | --- | --- |
| Parasite | Experiment | No of midguts | Prevalence (%) | Arithmetic mean | Median | Parasite range | P value |
| *c507* | Pool | 60 | 83 | 162.3 | 186 | 0-389 |  |
| *Δsto* |  | 77 | 22 | 0.7 | 0 | 0-11 | <0.0001 |
| *c507* |  | 80 | 78 | 56.0 | 19 | 0-558 |  |
| *Δrov* |  | 89 | 24 | 3.2 | 0 | 0-69 | <0.0001 |
| *c507* | R1 | 34 | 82 | 161.8 | 178 | 0-370 |  |
| *Δsto* |  | 43 | 21 | 0.6 | 0 | 0-7 | <0.0001 |
| *c507* |  | 15 | 80 | 52.3 | 19 | 0-193 |  |
| *Δrov* |  | 23 | 4 | 0.1 | 0 | 0-2 | <0.0001 |
| *c507* | R2 | 26 | 85 | 163.0 | 188 | 0-389 |  |
| *Δsto* |  | 34 | 24 | 0.9 | 0 | 0-11 | <0.0001 |
| *c507* |  | 30 | 73 | 65.2 | 19 | 0-533 |  |
| *Δrov* |  | 30 | 3 | 0.1 | 0 | 0 | <0.0001 |
| *c507* | R3 | 35 | 83 | 49.7 | 13 | 0-558 |  |
| *Δrov* |  | 36 | 53 | 7.8 | 1 | 0-69 | 0.0006 |
| Numbers of melanised parasites detected in the midguts of *c507* infected *CTL4* kd *A. coluzzii* mosquitoes at 7days post feeding. The P value was calculated using the Mann-Whitney U test. Data from pooled and independent biological replicates are presented. | | | | | | | |

| **Table S7. Sporozoite numbers and mosquito-to-mouse transmission after ookinete injection** | | | |
| --- | --- | --- | --- |
|  | Salivary gland sporozoites | | Infectivity to mice |
| Parasite | Mean | SEM |  |
| *c507* | 3,353(3,458; 3,248) | 74 | 4/4(2/2;2/2) |
| *Δsto* | 3,093(3,199; 2,987) | 75 | 4/4(2/2;2/2) |
| *c507* | 3,258(3,036; 1,888; 4,850) | 704 | 6/6(2/2;2/2;2/2) |
| *Δrov* | 2,439(2,960; 958; 3,400) | 614 | 6/6(2/2;2/2;2/2) |
| Mean salivary gland sporozoites at 21 days post *A. coluzzii* haemocoel inoculation with ookinetes, obtained from 6 biological replicates. Data from independent biological replicates are presented in brackets. Infectivity of sporozoites was assessed by infected mosquito bite back experiments of C57/BL6 mice at day 21 post haemocoel inoculation. Parasitaemia was monitored for 14 days post mosquito bite. SEM represents the standard error of the mean and ND represents not determined. | | | |

| **Table S8. Barcode and indexing primers** | | |
| --- | --- | --- |
| **Primer name** | **Sequence (5′ to 3′)** | **Primer description** |
| BC_STM Primer 91_Illumina | TCGGCATTCCTGCTGAACCGCTCTTCCGATCTGTAATTCGTGCGCGTCAG | Barcode upstream |
| BC_STM Primer 97_Illumina | ACACTCTTTCCCTACACGACGCTCTTCCGATCTCCTTCAATTTCGATGGGTAC | Barcode downstream |
| PE 1.0 Primer_Illumina PCR | **AATGATACGGCGACCACCGAGATCTACAC**TCTTTCCCTACACGACGCTCTTCCGATC*T | Index upstream |
| iPCRindex1 | **CAAGCAGAAGACGGCATACGAGAT**TGCTAATCACTGAGATCGGTCTCGGCATTCCTGCTGAACCGCTCTTCCGATC*T | Index 1 downstream |
| iPCRindex2 | **CAAGCAGAAGACGGCATACGAGAT**TAGGGGGATTCGAGATCGGTCTCGGCATTCCTGCTGAACCGCTCTTCCGATC*T | Index 2 downstream |
| iPCRindex3 | **CAAGCAGAAGACGGCATACGAGAT**AGTTTCCCAGGGAGATCGGTCTCGGCATTCCTGCTGAACCGCTCTTCCGATC*T | Index 3 downstream |
| iPCRindex4 | **CAAGCAGAAGACGGCATACGAGAT**CCTGGGAGGTAGAGATCGGTCTCGGCATTCCTGCTGAACCGCTCTTCCGATC*T | Index 4 downstream |
| iPCRindex5 | **CAAGCAGAAGACGGCATACGAGAT**ATACCACAAATGAGATCGGTCTCGGCATTCCTGCTGAACCGCTCTTCCGATC*T | Index 5 downstream |
| iPCRindex6 | **CAAGCAGAAGACGGCATACGAGAT**GATCTCTCGGGGAGATCGGTCTCGGCATTCCTGCTGAACCGCTCTTCCGATC*T | Index 6 downstream |
| iPCRindex7 | **CAAGCAGAAGACGGCATACGAGAT**ACCCTATACTCGAGATCGGTCTCGGCATTCCTGCTGAACCGCTCTTCCGATC*T | Index 7 downstream |
| iPCRindex8 | **CAAGCAGAAGACGGCATACGAGAT**CTCAATTAAGAGAGATCGGTCTCGGCATTCCTGCTGAACCGCTCTTCCGATC*T | Index 8 downstream |
| iPCRindex9 | **CAAGCAGAAGACGGCATACGAGAT**CGACAGAACGTGAGATCGGTCTCGGCATTCCTGCTGAACCGCTCTTCCGATC*T | Index 9 downstream |
| iPCRindex10 | **CAAGCAGAAGACGGCATACGAGAT**TCGCCATTATGGAGATCGGTCTCGGCATTCCTGCTGAACCGCTCTTCCGATC*T | Index 10 downstream |
| Primers used for the addition by overlap PCR of Illumina adaptors and indices to the samples of pools of transgenic parasites collected. Highlighted are, in green-barcode binding sites; bold-Illumina adaptors and underlined- i7 indices. All primers are listed in a 5′ to 3′ direction. | | |

| **Table S9. Primers for generation of transgenic parasites and protein expression** | | |
| --- | --- | --- |
| **Primer name** | **Sequence (5′ to 3′)** | **Description** |
| P1 F | TTGGGCCCGTATATTGCATGCTATTCAATTGTATTG | CRONE disruption upstream target ApaI |
| P2 R | CCAAGCTTGAACTTTTCCTTTGGTTATGCAATAATG | CRONE disruption upstream target HindIII |
| P3 F | TGAATTCGATGATCAACCAAATTTTTTGTAGAG | CRONE disruption downstream target EcoRI |
| P4 R | TTGGATCCGCGCATAAGTGCACACTTGTTATATAG | CRONE disruption downstream target BamHI |
| P5 F | TTGGGCCCGTGCACTAAGATTTGCGCATTTTATCAAATG | ROVER disruption upstream target ApaI |
| P6 R | CCAAGCTTCAAAAGATTATAAAATTAATAAGTC | ROVER disruption upstream target HindIII |
| P7 F | TGAATTCGAATATTAAAAAATTAGCTATAATTAG | ROVER disruption downstream target EcoRI |
| P8 R | TTGGATCCCACACATATGTGTGTATATCCAATATTC | ROVER disruption downstream target BamHI |
| P9 F | TTGGGCCCGTTGCTGTATCTACACATTTTAACCTG | SPM1 disruption upstream target ApaI |
| P10 R | CCAAGCTTGGAAAGTCGTATATTCTTTTCGTTTTATG | SPM1 disruption upstream target HindIII |
| P11 F | TGAATTCGTGCAGAGATGTATGAGTACATAGATGT | SPM1 disruption downstream target EcoRI |
| P12 R | TTGGATCCGCATAATTGGATATATATACTTGG | SPM1 disruption downstream target BamHI |
| PlasmoGEM HAP2 GT R (P13) | TGCCCCATTTATTTTTGTCCT | Diagnostic primer WT and KO |
| PlasmoGEM GW2 (P14) | CTTTGGTGACAGATACTAC | Diagnostic primer WT |
| PlasmoGEM HAP2 QCR1 (P15) | TGCAGATACATCTCCGTCAGGT | Diagnostic primer WT |
| PlasmoGEM HAP2 QCR2 (P16) | TGTTGTGTTTCCTCCATCCA | Diagnostic primer KO/c507 |
| PlasmoGEM STONES GT F (P17) | TGGACCCGAGGATGTCACATGGG | Diagnostic primer WT and KO/c507 |
| PlasmoGEM STONES QCR1 (P18) | TGCGCATTCTGCTCCTGGGG | Diagnostic primer WT and KO/c507 |
| PlasmoGEM STONES QCR2 (P19) | CACAGTTTGTGCAGAGAAT | Diagnostic primer WT/c507 |
| PlasmoGEM CRYSP GT R (P20) | ATTGCACGTTGAATATGCCA | Diagnostic primer WT and KO/c507 |
| PlasmoGEM CRYSP QCR1 (P21) | CGAGGCGGAATGCAAAAAGCT | Diagnostic primer WT/c507 |
| PlasmoGEM CRYSP QCR2 (P22) | CCACTTCCCAAACAGAGGATGTAGCC | Diagnostic primer KO/c507 |
| CRONE INT F (P23) | GCATTGTTTATATACGCTTCATAAGTTTTG | Diagnostic primer WT and KO/c507 |
| CRONE WT R (P24) | CATAAGGCAATTTTTTAACTAAGGCCAC | Diagnostic primer WT/c507 |
| ROVER INT F (P25) | GCTTATGTTATTATTTAATATCCCCTT | Diagnostic primer WT and KO/c507 |
| ROVER WT R (P26) | CCATTTCCACACATAACTGAATTTGTTCC | Diagnostic primer WT/c507 |
| SPM1 INT F (P27) | GGTTTATGGACAAAAAAACAAATTAGCAG | Diagnostic primer WT and KO/c507 |
| SPM1 WT R (P28) | CTTCATCAAAAAAATTAAAATTTACAC | Diagnostic primer WT/c507 |
| TgDHFR 5'UTR R (P29) | GATGTGTTATGTGATTAATTCATACAC | Diagnostic primer KO/c507 |
| ROVERHA 5’ F (P30) | GATTACGCCAAGCTTGGGCCCGAATGACCCTAGATACAATGATTATCCACCTATGAACGGATTTGG | ROVER tagging upstream target ApaI |
| ROVERHA 5’ R (P31) | CTAAGCATAGTCTGGAACGTCATAAGGGTATGCGTAATCTGGCACGTCGTATGGATATGCA | ROVER tagging upstream target |
|  | TAATCTGGTACATCGTATGGGTAATTTCCTACACCAATATTTAAGTCAATTATAACTCC |  |
| ROVERHA 3’ UTR F (P32) | GACGTTCCAGACTATGCTTAGGCAAAAGATTATAAAATTAATAAGTCAAGAGTTTG | ROVER 3’UTR |
| ROVERHA 3’ UTR R (P33) | CTAGAGCGGCCGCCACCGCGGGTGCACTAAGATTTGCGCATTTTATCAAATGTTTACC | ROVER 3’UTR SacII |
| ROVERHA 3’ F (P34) | CTTCAATTTCGGGTACCCTCGAGGCAAAAGATTATAAAATTAATAAGTCAAGAGTTTG | ROVER tagging downstream target XhoI |
| ROVERHA 3’ R (P35) | GAATTCGCGGCCGCCCCGGGGGCGATAACAAGAATGATGATAATAACGATCCAGGAG | ROVER tagging downstream target XmaI |
| PlasmoGEM STONESHA GT R (P36) | TGCCAAACTGTCAGAGGCATCA | Diagnostic primer tag |
| PlasmoGEM STONESHA QCR1 (P37) | AGACCATTACATGCCGCAAA | Diagnostic primer tag |
| PlasmoGEM STONESHA QCR2 (P38) | ACAAAACGGCAACTGACTTTGC | Diagnostic primer tag |
| PlasmoGEM CRYSPHA GT F (P39) | TCCCCACCTTTTCCTTCCCT | Diagnostic primer tag |
| PlasmoGEM GW1 (P40) | CATACTAGCCATTTTATGTG | Diagnostic primer tag |
| PlasmoGEM CRYSPHA QCR1 (P41) | GGCTACATCCTCTGTTTGGGAAGTGG | Diagnostic primer tag |
| PlasmoGEM CRYSPHA QCR2 (P42) | ACCAGCACTCGGAATTGCCA | Diagnostic primer tag |
| PlasmoGEM CRONEHA GT F (P43) | AGTGGGGGAAGATGGGGAGGA | Diagnostic primer tag |
| PlasmoGEM CRONEHA QCR1 (P44) | CCGTCCACACAAACGGCCCA | Diagnostic primer tag |
| PlasmoGEM CRONEHA QCR2 (P45) | TGTTTGCTTTTATTAAGAATGGGCT | Diagnostic primer tag |
| ROVERHA INT F (P46) | GCAGATTCACATAACGAAACTCATAATATATTG | Diagnostic primer tag |
| HA INT R (P47) | GTATGCGTAATCTGGCACGTCGTATG | Diagnostic primer tag |
| ROVERHA WT R (P48) | GATCTATTATTAAACCATATTCATTTAG | Diagnostic primer tag |
| CRONE GA F (P49) | AGCTCCGTCGACAAGCTTGCGGCCAAGATCCCCATCGAGAACCCCCC | STONE optimized expression primer |
| CRONE GA R (P50) | TGGTGGTGGTGCTCGAGTGCGGCCGCCTGAGCGGTCTGGGTGGAGG | STONE optimized expression primer |
| CTL4F (P51) | TAATACGACTCACTATAGGGTGGTTTGATGCCGTGTCCT |  |
| CTL4R (P52) | TAATACGACTCACTATAGGGAATAAATTGTCTCGGTTCATCATC |  |
| PlasmoGEM MAP2 GT F (P53) | ACCATGAGTGCATGCATAGGA | Diagnostic primer WT and KO |
| PlasmoGEM MAP2 QCR1 (P54) | ACGAATCACAATTGACCAGGCT | Diagnostic primer WT |
| PlasmoGEM MAP2 QCR2 (P55) | TGCGTGTTTGTGAAACTAATGAGGCA | Diagnostic primer KO/c507 |
| Where appropriate, target restriction sites are shown as underlined italics. The appropriate restriction enzyme is presented in the description column. F, forward; R, reverse; INT, integration; WT, wild-type; KO, knockout; UTR, untranslated region; GA, Gibson assembly. All primers are listed in a 5′ to 3′ direction. | | |
